# Programming protein shape as an explicit design layer via CAD blueprint-guided diffusion

**DOI:** 10.64898/2026.07.22.740177

**Authors:** Yilun Qi, Guoxun Zhang, Klaus Yserentant, Sonya Lee, Artem Lyubimov, Kyrellos Ibrahim, Giuseppe Cimicata, James S. Fraser, Bo Huang, Tanja Kortemme

**Affiliations:** Department of Bioengineering and Therapeutic Sciences, University of California, San Francisco; San Francisco, CA, USA; Department of Pharmaceutical Chemistry, University of California, San Francisco; San Francisco, CA, USA; Quantitative Biosciences Institute (QBI), University of California, San Francisco; San Francisco, CA, USA; Department of Biochemistry and Biophysics, University of California, San Francisco; San Francisco, CA, USA; Biohub San Francisco, San Francisco, CA, USA

## Abstract

In engineering, industrial design principles link geometric form with function. In nature, global protein shape, curvature, and chirality similarly drive critical processes like membrane remodeling and cellular motility. However, explicitly programming these global features remains a challenge for generative AI. Here, we integrate computer-aided design (CAD) blueprints with diffusion models, establishing protein shape as an explicit, programmable layer. We engineered diverse, tunable architectures, including single-chain shapes, scaffolds with unusual twist handedness, and superhelical assemblies, structurally validating 31 of 59 designs via X-ray crystallography or electron microscopy. To demonstrate shape-encoded function, we coupled fluid-flow-optimized helical propellers to an F_1_-ATPase rotor, yielding a prototype ATP-dependent molecular swimmer. This framework translates industrial design principles to the molecular scale, enabling the programmable engineering of biomimetic architecture with emergent functions.

**One-sentence summary:** Integrating CAD blueprints with diffusion-based AI translates industrial design principles to the molecular scale, enabling the explicit programming of protein shapes with emergent functions.

## Main Text

Protein function is governed by both global architecture and local structural features. Much of the field of computational design of new protein functions has focused on local features: Atomic-scale placements of amino acid side chains in protein-protein interfaces and catalytic sites can now be designed with considerable accuracy, enabling exciting advances in generating protein-protein binders and enzymes (*1*, *2*). However, for many biological processes, global three-dimensional shape and higher-order assembly architectures are equally critical determinants of function (*3–6*). Shape governs biological behaviors across scales: single-chain repeat proteins can form elongated, curved, or solenoid-like shapes that create extended interaction surfaces (*7*) and the toroidal DNA sliding clamps encircle DNA to enhance polymerase processivity (*6*, *8*). At higher-order scales, the specific curvature of protein assemblies is important for membrane remodeling (*9*, *10*), the superhelical filaments of flagella drive motility (*11*, *12*), and large-scale needle-like structures are used to inject bacterial pathogens into host cells (*13*).

Despite this importance, protein shape remains difficult to access as a fully programmable design layer. For single-chain proteins, diffusion-based generative models can in principle condition on approximate global shapes but have not been experimentally validated directly (*14–16*), or constraints have been imposed as soft matching during sampling rather than as continuously enforced shape potential, requiring downstream filtering or refinement to improve agreement with the target shape (*17*). In parallel, reinforcement-learning approaches based on Monte Carlo tree search have sampled and connected local helical elements within prescribed shapes (*18*), but the discreteness of the sampled elements can limit shape completeness, accessible diversity, and programmability. Similarly, symmetry-based strategies have led to exciting advances in machine (*19*) and vaccine design (*20*, *21*), but their reliance on idealized or standardized building blocks restricts component shape diversity, and the prescribed symmetry provides only relatively coarse control over the final assembly envelope (*22–24*). More recent modular and deep learning-based approaches have expanded the range of accessible protein structures and assemblies (*25*), but these methods also specify motifs, interfaces, block connectivity, or symmetry relationships rather than continuously enforcing an independently defined external shape envelope. Thus, global protein shape is typically an indirect outcome of symmetry, block choice, interface design, or model capabilities and existing approaches lack the programmability of generalizable computer-aided design (CAD) frameworks for specifying and tuning protein assembly shapes that ultimately enable physical behaviors found in nature such as fluid transport, motion, or mechanical response.

Here we introduce a diffusion-based framework for programmable control of protein shape through CAD-guided shape blueprinting. We reasoned that by incorporating CAD models for explicit specification of a target shape, our approach should be generalizable as well as broadly and easily accessible. We furthermore sought to develop a shape potential that could be continuously enforced during structure generation to maximize agreement with user-defined shapes, while keeping interoperability with existing generative protein design workflows using diffusion models. To experimentally validate the approach, we pursued design goals of increasing difficulty and scale: We constructed (i) proteins with CAD-defined non-canonical external shapes, (ii) scaffolds with common topology but different twist handedness, and (iii) shared protein building blocks with controllable external geometries. Extending scale, we (iv) constructed higher-order architectures where tunable global shape, curvature, chirality, and organization arise directly from tuning the local shape states of the building blocks as a programmable parameter. Finally, to link these tunable protein architectures to dynamic function, we used hydrodynamic simulations to predict and build assemblies with specified fluid flow properties and created prototype ATP-dependent swimmers by linking these designed assemblies to an F₁-ATPase rotor.

### CAD-guided blueprinting for programmable protein shape

To explicitly program protein shape, we sought to develop a method that incorporates CAD-defined shape blueprints into a diffusion-based deep-leaning backbone design framework (**Fig. 1**); here we implemented the method in the widely used RFdiffusion model. Input blueprints can be readily generated using standard CAD software or sourced from public 3D repositories. During diffusion, the evolving protein backbone is represented as a Cα point cloud and continuously guided toward the target shape envelope by the CAD-derived shape potential (**Fig. 1A**). Our guiding potential combines two complementary constraints: a boundary term that discourages points from leaving the blueprint-defined volume, and a global shape-matching term that promotes agreement with the target shape. We implemented these constraints using signed distance functions and point-cloud similarity metrics (Methods). Together, these constraints discourage solutions that escape the target volume or poorly agree with the global shape blueprint. As an example, denoising trajectories starting from Gaussian noise progressively converge toward a predefined conical shape during diffusion, ultimately producing protein backbones that followed the prescribed blueprint envelope (**Fig. 1B**).

**Fig. 1.**
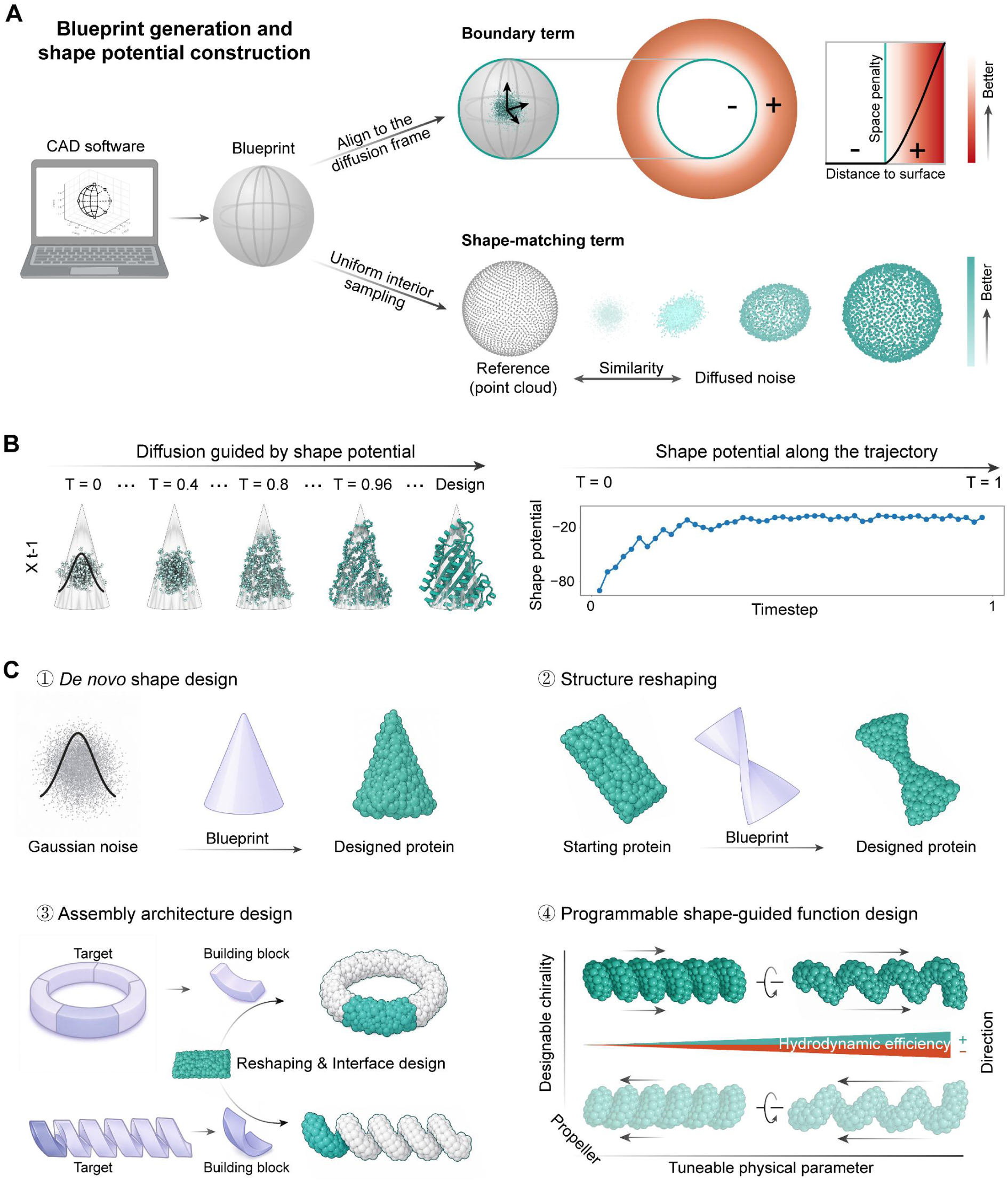
CAD-defined shape potential enables programmable control of protein shape. (A) Schematic of the blueprint-derived shape potential, which combines a boundary term (upper right) and a shape-matching term (lower right). The boundary term uses a signed distance function (SDF) to penalize residues located outside the CAD-defined shape envelope, thereby enforcing spatial confinement. The shape-matching term uniformly samples the CAD blueprint to generate a reference point cloud and compares it with the evolving protein backbone, represented as a Cα point cloud, using the Chamfer distance (Methods) to promote global shape agreement. (B) Shape-conditioned diffusion trajectory showing progressive convergence of the backbone from noise toward a target conical shape, accompanied by convergence of the shape potential over diffusion timesteps. (C) Conceptual overview schematic illustrating the shape-conditioned design capabilities enabled by the CAD-defined shape potential, including 1. de novo shape design, 2. topology-preserving structure reshaping, 3. assembly architecture design, and 4. programmable shape-guided function design.

More broadly, we reasoned that this framework could support multiple modes of shape-conditioned protein design, all of which we validate experimentally in this study: In the first mode, *de novo* backbones can be generated from Gaussian noise to conform to specific blueprints (**Fig. 1C**, panel 1). In the second, existing protein backbones can be twisted and reshaped while preserving the fold topology (**Fig. 1C**, panel 2). In the third, by tuning the local shape states of shared building blocks, the framework can be extended to the assembly scale, enabling construction of higher-order protein architectures with programmable physical parameters such as curvature, diameter, helical pitch, and chirality (**Fig. 1C**, panel 3). Finally, these programmable architectures provide a basis for exploring shape-encoded physical properties, such as tunable hydrodynamic behaviors (**Fig. 1C**, panel 4).

### *De novo* generation of proteins with prescribed shape envelopes

To test whether shape-conditioned diffusion can generate proteins with prescribed external shape envelopes (**Fig. 1C**, panel 1), we used five CAD blueprints as target shapes (**Fig. 2A**). We chose these blueprints for features such as a 90-degree turn, sharp boundaries, and specific surface curvature. For each target, we generated 100–200 candidate protein backbones and selected 20–30 representative backbones with the strongest agreement with the expected shape for sequence design (**Fig. 2B**, **Fig. S1A, B**).

**Fig. 2.**
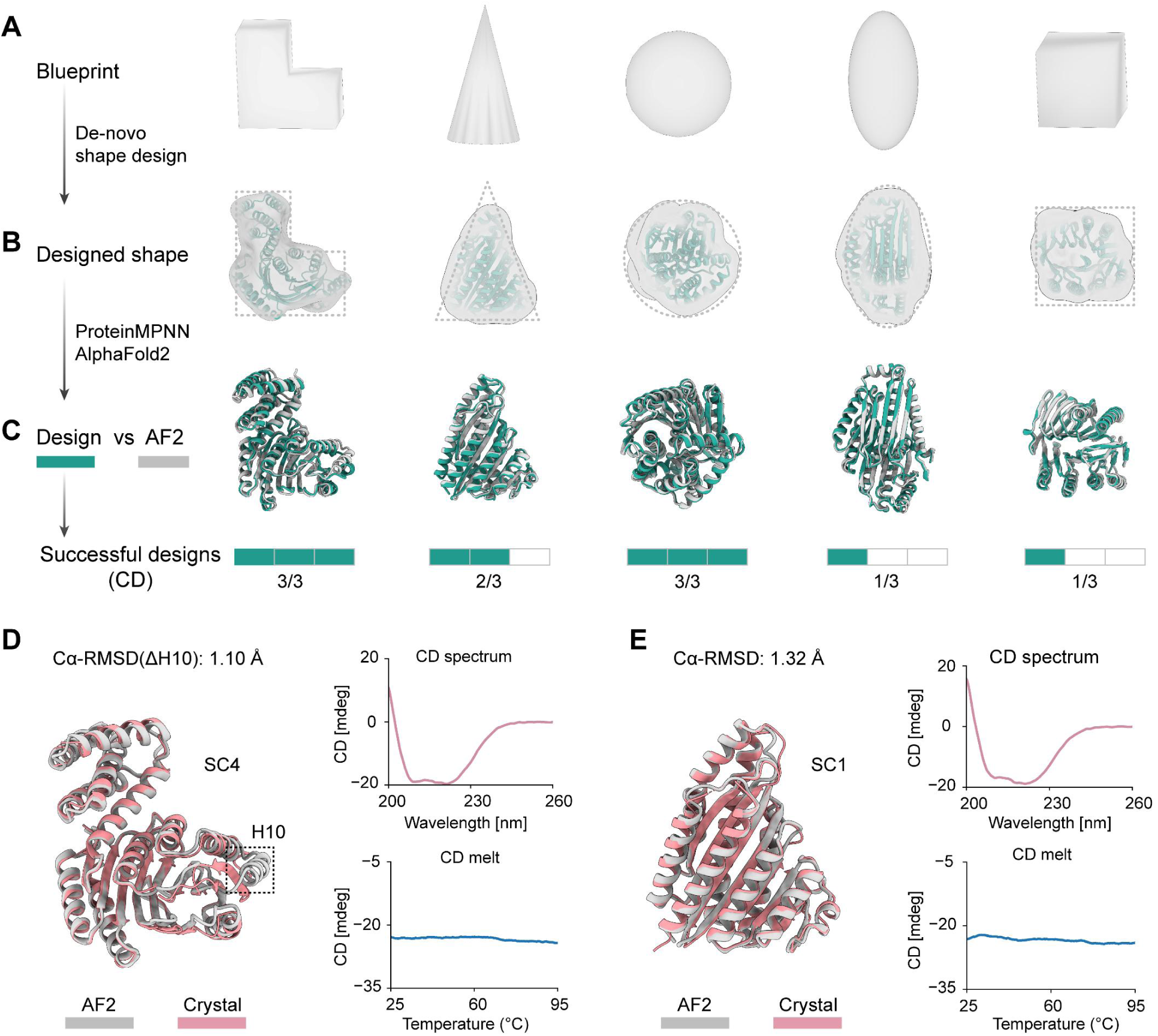
*De novo* shape-conditioned diffusion generates proteins that recapitulate prescribed three-dimensional shape envelopes. (A) CAD blueprints defining the target shape envelopes used for *de novo* design, shown from left to right: bracket, cone, sphere, ellipsoid, and cube. (B) Representative backbone models conforming to the target envelopes. Designed structures are shown in green, and the corresponding blueprint outlines are shown as dashed lines. (C) Structural agreement between designed models, shown in green, and AlphaFold2 predictions, AF2, shown in gray, after sequence design with ProteinMPNN. Three designs were ordered for each target shape. Numbers of successful designs with stable secondary structure as measured by circular dichroism (CD) are shown below each design target shape. (D) Biophysical validation of the bracket design SC4. Superposition of the AF2 model, gray, and the crystal structure, pink, yielded a Cα RMSD of 1.01 Å after excluding a locally rearranged helix (H10), boxed. CD spectrum and thermal denaturation indicate a well-folded and thermally stable α/β structure. (E) Biophysical validation of the cone design SC1. Superposition of the AF2 model, gray, and the crystal structure, pink, yielded a Cα RMSD of 0.90 Å. CD spectrum and thermal denaturation indicate a well-folded and thermally stable α/β structure.

We next examined whether these shape-constrained backbones retained sufficient sequence designability to support stable folded structures. We designed eight sequences for each backbone with ProteinMPNN and evaluated self-consistency with the design model by predicting their structures with Alphafold2 and comparing the result to the design model (**Fig. 2B, C** and Methods). Cone, sphere, and ellipsoid designs showed higher *in silico* success rates (AF2 predicted structures matching the design model with < 2.0 Å Cα RMSD) than the bracket and cube targets (**Fig. S1C**); designability decreased with increasing chain length (**Fig. S1D**). These trends are consistent with the greater difficulty of encoding sharp turns and corners within protein folds and of generating designable long-chain proteins. We then selected 15 designs for experimental validation based on Cα RMSD (<1.5 Å) and pLDDT (> 90). Of these 15 designs, 11 were solubly expressed in *E. coli*, and 10 produced a dominant size-exclusion chromatography peak at retention volumes consistent with the expected target monomer size (**Fig. S2A**). Circular dichroism spectroscopy and thermal denaturation assays further indicated that these proteins were well folded and thermally stable (**Fig. 2D, E**; **Fig. S2B**).

To experimentally assess whether the designs adopted the predefined shapes, we determined crystal structures of two designs, Bracket SC4 and Cone SC1, at resolutions of 1.72 Å and 2.70 Å, respectively (**Table S1**). The SC1 structure closely matched the design model, with a backbone Cα RMSD of 1.32 Å, while preserving the intended conical shape envelope (**Fig. 2E**). The SC4 structure also recapitulated the designed bracket-shaped external contour, although one local fragment, residues 214-234, corresponding to helix 10 (H10) in the design, rearranged from the designed helix into a β-strand (**Fig. 2D**; **Fig. S2C**) with a backbone Cα RMSD to the design model of 2.39 Å. Excluding this local rearrangement, the backbone Cα RMSD was 1.10 Å, and the overall shape envelope remained intact. Together, these results show that shape-conditioned diffusion can generate experimentally folded proteins with prescribed external shape envelopes, establishing global protein shape as an explicit designable variable in *de novo* protein design.

### Reshaping protein backbones into alternative shape states

We next evaluated whether shape-conditioned diffusion can reshape existing protein backbones towards prescribed global shape states (**Fig. 1C**, panel 2). In particular, we sought to reshape α/β proteins containing extended β-sheet layers to adopt a left-handed twist, a rare structural state found in only a minor fraction of naturally occurring β-sheet proteins, as opposed to the dominant right-handed twist (*26*, *27*). We first constructed an α/β backbone model with defined topology (**Fig. S3A**). To diversify backbone architecture and chain length, we performed fold-conditioned diffusion with variable loop insertions by RFdiffusion (**Fig. 3A**, **Fig. S3B**). 145 out of 200 retained the original 2-layer *α/β* backbones architecture after diffusion.

**Fig. 3.**
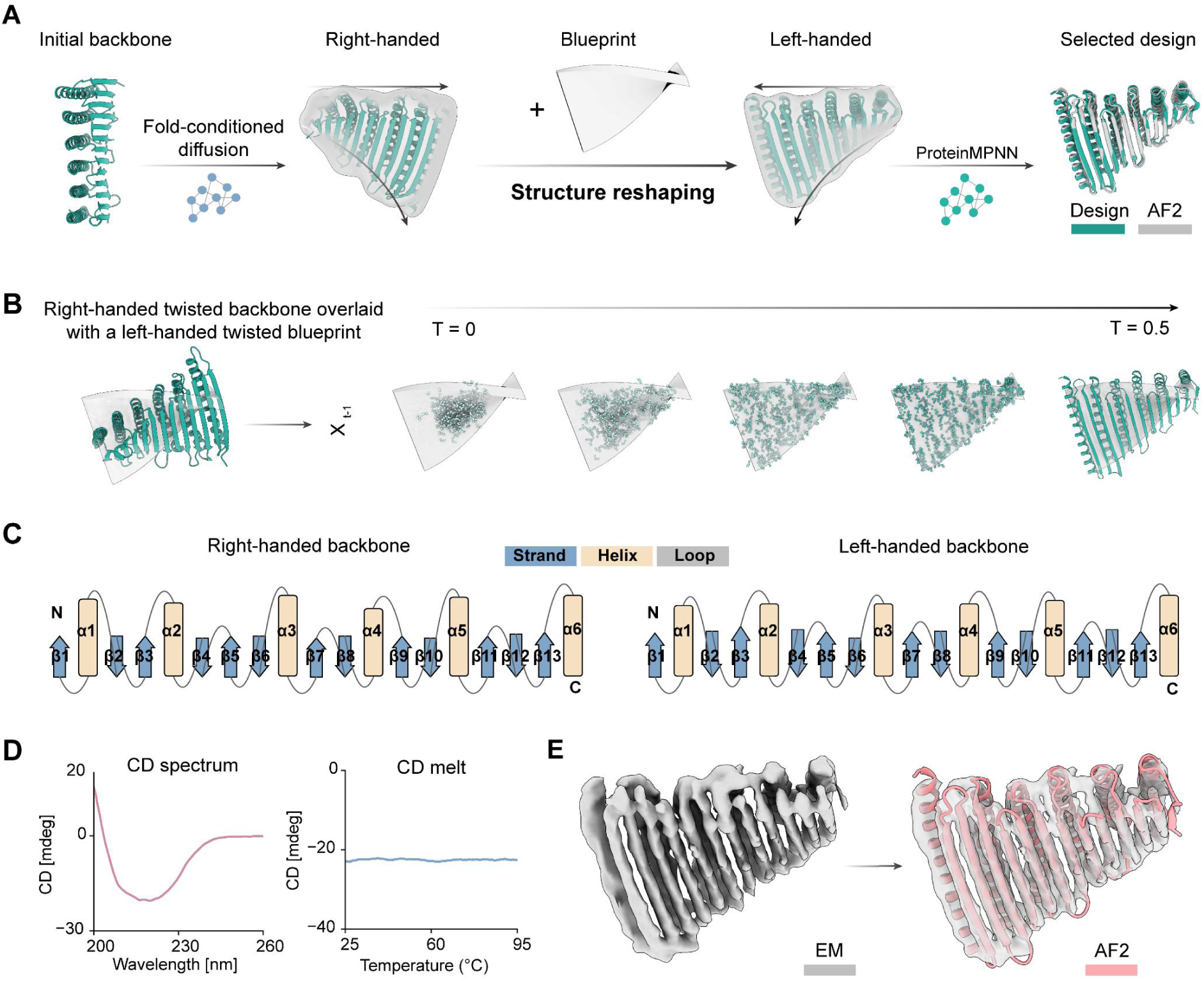
Shape-guided reshaping enables access to rare β-sheet twist chirality while preserving backbone topology. (A) Design strategy for topology-preserving structure reshaping. Fold-conditioned diffusion generates right-handed α/β backbones from an initial backbone model; 145 of 200 trajectories preserved a regular 2-layer architecture, and all resulting 145 models adopted the right-handed twist. CAD blueprints encoding left-handed twist were then applied to reshape these backbones, yielding candidates whose AF2 predicted structures (gray) after ProteinMPNN sequence design agreed with the designed backbones (green). (B) Representative reshaping trajectory. A right-handed twist backbone was placed within a left-handed blueprint, and diffusion progressively transformed the structure toward the target twist chirality. (C) Topology schematics of the backbone retained through structure reshaping showing small changes in secondary structure lengths. (D) Circular dichroism spectrum and thermal denaturation indicate that the selected design adopts a well-folded and thermally stable α/β structure. (E) Cryo-EM reconstruction (gray) of the selected design reveals a β-sheet architecture consistent with the AF2 model (pink).

We next imposed CAD blueprints encoding left-handed twist to reshape these right-handed twist backbones (**Fig. 3A**). To sample diverse alignments between the starting right-handed twist backbones and target shape envelopes, we generated multiple blueprint orientations (**Fig. S3C**). Shape-conditioned diffusion redirected a subset of trajectories in which the starting backbone was productively aligned with the blueprint toward left-handed β-sheet architectures. A representative trajectory shown in **Fig. 3B** illustrates progressive reshaping towards the prescribed left-handed state during diffusion, accompanied by corresponding changes in the shape potential **(Fig. S3D**). Comparison of structures before and after reshaping showed that the order and connectivity of secondary-structure elements were preserved despite small length shifts of specific secondary-structure elements to conform to the specified global shape (**Fig. 3C**, **Fig. S3E**), indicating that a common fold topology can accommodate distinct external shape states. (In contrast, unconstrained fold-conditioned RFdiffusion sampling retained right-handed twist, indicating that the left-handed state was induced by the external shape constraint rather than by the generative model prior.) We next examined whether reshaped left-handed backbones retained sufficient sequence designability to support stable folding (Methods). Left- and right-handed backbones showed comparable *in silico* success rates after sequence design and AF2 prediction (**Fig. S3F**), indicating that reshaped left-handed backbones remained highly designable.

Six candidate sequences with strong agreement between AF2 predictions and the target rare left-handed backbones were selected for experimental characterization (**Fig. 3A**). The selected designs were expressed in *E. coli* and purified (Methods). One design showed soluble expression at preparative scale and produced a dominant size-exclusion chromatography peak suitable for structural characterization (**Fig. S4A**). Circular dichroism spectroscopy and thermal denaturation experiments confirmed that the protein was well folded and thermally stable (**Fig. 3D**). Although the purified sample eluted predominantly as a single peak by size-exclusion chromatography, cryo-EM classification revealed both monomeric and dimeric particle classes (**Fig. S4A, B**).

A 4.0 Å reconstruction of the monomeric class resolved the overall arrangement of secondary-structure elements and closely matched the AF2-predicted design model in global shape, curvature, and secondary-structure organization (**Fig. 3E**). Although we could not establish absolute handedness from the density alone, the reconstruction and model fit were consistent with the designed left-handed twist of the β-sheet layer (see Methods for details on evaluation of reconstruction handedness). Discontinuous density near the N-terminal β-sheet suggested local flexibility in this region (**Fig. S4C**). A substantial fraction of particles formed a dimeric class, yielding an additional 4.1 Å reconstruction that likewise matched the overall architecture and designed twist state (**Fig. S4B, D, E**). Despite local disorder near the N terminus, both reconstructions supported formation of the intended curved architecture. Together, these results demonstrate that shape-conditioned diffusion can reshape existing protein backbones into shape states that are strongly underrepresented in natural protein structures while largely preserving overall topology. These findings support the view that global protein shape can be specified as a design variable distinct from fold topology, enabling existing backbones to be redirected toward alternative shape states through externally imposed constraints.

### Encoding higher-order protein architectures through programmable local shape states

To test whether shape-conditioned diffusion could be extended to programmable assembly design (**Fig. 1C**, panel 3), we developed a framework for constructing target higher-order protein architectures from modular shape building blocks (**Fig. 4A**). CAD models of target architectures were first decomposed into repeating units, which served as blueprints for reshaping a shared backbone scaffold into distinct curvature and twist states. The reshaped monomers were then docked and assembled through designed intermolecular interfaces compatible with the target architecture. In this strategy, the local shape state of each building block functions as an architectural parameter that specifies the global organization of the assembly.

**Fig. 4.**
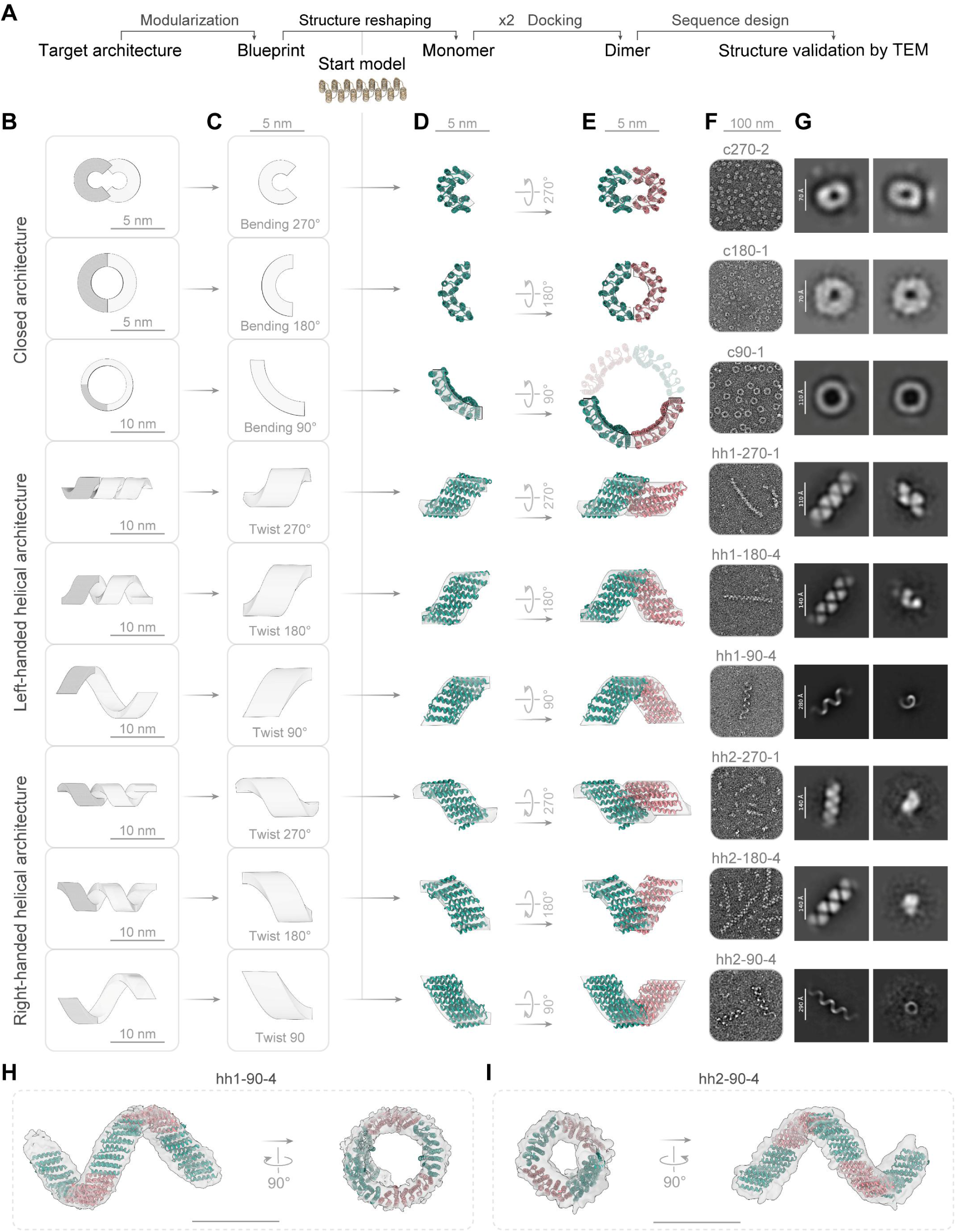
Programmable assembly of diverse protein architectures by tuning the shape of a shared building block. (A) Design approach. Starting from a target architecture, the assembly was first modularized into component building blocks, which were then used to define CAD blueprints for local shape states. A shared 340-residue helical bundle was then reshaped according to each blueprint to generate monomeric building blocks. Reshaped monomers were docked to form dimeric assembly units for interface design using ProteinMPNN, followed by structural validation using AF2 and experimental validation of the assembled architectures by negative-stain transmission electron microscopy, nsTEM. (B–E) Architecture design across closed assemblies, top, left-handed helical assemblies, middle, and right-handed helical assemblies, bottom. (B) Target architectures. (C) Corresponding blueprints specifying curvature or twist. (D) Reshaped monomeric backbones. (E) Dimeric building blocks generated by blueprint-guided docking. Panels C-E share the scale bars shown at the top. (F, G) Experimental validation by nsTEM. Representative particles are shown in (F) with scale bar at the top, and representative 2D class averages are shown in (G), confirming the designed architectures. In G, scale bars for target architectures are indicated individually. Class averages were derived from negative-stain TEM data, except for hh1-90-4 and hh2-90-4, which were derived from cryo-EM data. (H, I) Cryo-EM reconstructions validate the overall assembled architectures. Fitted models agree with the reconstructed densities for the helical assemblies designed to be left-handed (H) and right-handed (I).

We first evaluated three closed architectures constructed from a shared starting scaffold reshaped into distinct curvature states corresponding to 270°, 180° and 90° bending states (**Fig. 4B, C**, rows 1-3), denoted c270, c180 and c90, respectively. An ideal helical bundle served as the common starting scaffold and was reshaped into the distinct curved shapes (**Fig. 4D**, rows 1-3). Guided by these local shape states, 2 repeating units were docked to create the assembly interface. Interface-designed units assembled into closed architectures with distinct global organizations (**Fig. 4E**). Negative staining TEM (nsTEM) imaging and 2D class averages supported assembly into the intended closed structures, including elliptical and circular ring assemblies across the curvature series (**Fig. 4F, G**). Three-dimensional reconstructions matched the overall shapes of the corresponding design models (**Fig. S6A**). These results show that control over local building-block curvature can be propagated into programmable closed protein architectures.

We next asked whether this framework could be expanded to open and extendable architectures. We selected helical architectures with varying pitch and diameter as test cases (**Fig. 4B**, rows 4-6). Corresponding building blocks were derived from blueprint-guided reshaping of the same initial scaffold, now incorporating both curvature and axial twist (**Fig. 4C, D**). Designs with twist states of 270°, 180° and 90° were denoted hh1-270, hh1-180 and hh1-90. Experimental characterization showed assembly into helical filaments across the architecture series (**Fig. S6B**). nsTEM images and 2D class averages revealed progressively more open assemblies across the programmed shape-state series (**Fig. 4F, G**). Three-dimensional reconstructions of representative designs closely matched the intended helical shapes (**Fig. 4H**; **Fig. S6B**), indicating that systematic tuning of local shape states enables control over emergent higher-order architecture with different physical parameters.

We further tested whether superhelical handedness could be programmably inverted using the same backbone scaffold and reshaping strategy (**Fig. 4B**, rows 7–9). By introducing CAD blueprints encoding helical architectures with reversed handedness, we generated the right-handed counterparts hh2-270, hh2-180, and hh2-90. Electron microscopy and three-dimensional reconstructions yielded filament architectures that closely matched the corresponding design models in overall morphology and helical geometry. Although the 7-10 Å reconstructions did not independently determine absolute handedness, their agreement with the predicted structures supported realization of the designed inversion of superhelical geometry (**Fig. 4I**; **Fig. S6C**). These results support programmable control of global assembly handedness through local shape states encoded in shared building blocks.

Together, these results show that assembly architecture can be programmed through the shape of its constituent building blocks. This differs from approaches that specify only an overall assembly envelope or primarily optimize predefined symmetric interfaces, because in our approach the local curvature and twist of the shared scaffold are designed as architectural parameters. By matching building-block shape to the target assembly geometry, this strategy enabled closed, open, and handedness-inverted assemblies, including smooth, extensible superhelical coil architectures produced by the continuous propagation of designed block curvature. Such architectures remain difficult to generate using conventional assembly design strategies, highlighting high-order architecture as an explicit design layer for *de novo* protein assembly design.

### Shape-programmed architectures couple protein shape to emergent physical function

To test whether blueprint-guided shape design could extend beyond programmable architecture to emergent physical behavior, we performed low-Reynolds-number hydrodynamic simulations on representative protein assemblies generated through the programmable local shape states described above. The simulated architectures were grouped as symmetric ring controls, referred to here as closed-ring assemblies and tubular helical propellers, referred to as helical assemblies. Using rigid-body rotational simulations (Methods), we evaluated whether these architectures could convert rotational motion into directional flow fields and net axial flux. Each structure was subjected to identical rotational driving, and the resulting velocity fields and axial transport profiles were computed **(Fig. S7A–J**).

Representative simulations revealed that directional pumping depends on global architecture and handedness (**Fig. 5A**). As expected, the symmetric ring control produced negligible net axial pumping. By contrast, left- and right-handed helical propellers generated coherent axial flows of opposite sign under the same rotational convention. Thus, shape-programmed chirality provides a mechanism for breaking hydrodynamic symmetry and defining the direction of axial transport, consistent with classical symmetry-breaking principles at low Reynolds number (*28*). Notably, a rotating F-actin control assembly with a comparable axial extent also produced negligible net pumping (**Fig. 5A**, **Fig. S7B**), despite its intrinsically chiral filament organization. This result confirms that efficient directional transport does not only arise from symmetry-breaking by chirality, but also depends on the coupling between local building-block shape and the global assembly architecture.

**Fig. 5.**
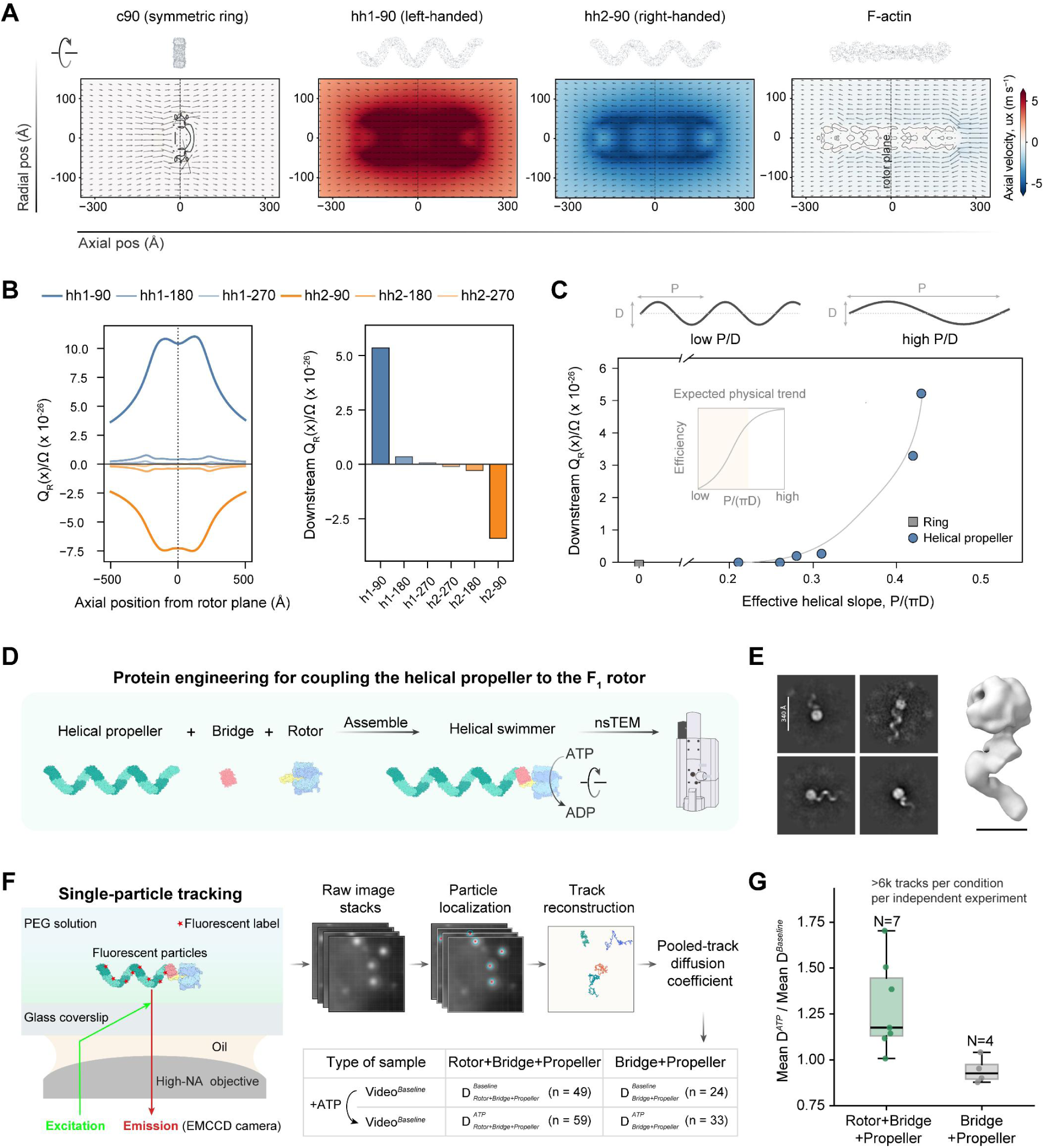
Shape-programmed architectures encode emergent physical functions. (A) Stokes-flow simulations of representative rotating structures: a symmetric ring, c90; a left-handed helical propeller, hh1-90; a right-handed helical propeller, hh2-90; and F-actin. Color scale indicates the axial velocity component, (u_x), and arrows indicate in-plane velocity vectors in the axial-radial section. Opposite-handed helical propellers generated axial flows of opposite sign, whereas the symmetric ring produced negligible net pumping. (B) Architecture-dependent pumping in the helical propeller series. Left, finite-disk sectional axial flux, ***Q_R_*(*x*)/Ω**, along the rotation axis. Right, downstream averaged ***Q_R_*(*x*)/Ω** for each helical propeller design. Pumping strength and direction varied with helical architecture, with hh1-90 and hh2-90 producing the strongest flows in opposite directions. (C) Architecture-function landscape relating downstream pumping to the effective helical slope, ***P***/(***π***), for helical propellers, where *(P)* is the pitch and *(D)* is the characteristic diameter. Across the helical propeller series, downstream pumping increased with effective helical slope, consistent with the expected physical trend for low-Reynolds-number helical pumping (inset). ***Q_R_*(*x*)** denotes the local finite-disk sectional axial flux rather than conserved flux across an infinite cross-section. (D) *In vitro* assembly. Purified helical propeller, bridge protein, and F₁-ATPase rotor were sequentially assembled for structural and functional characterization. (E) Representative 2D class averages and ab initio 3D reconstruction from negative-stain TEM data showing assembled particles consistent with the designed Rotor-Bridge-Propeller architecture. (F) Schematic of the single-particle tracking assay used for functional characterization of assembled complexes, where n in the table denotes the number of videos collected. (G) Ratio of the mean diffusion coefficient after ATP addition to that before ATP addition. N denotes the number of independent experiments. Each point represents one independent experiment comprising at least three videos before and after ATP addition (Video*^Baseline^* and Video*^ATP^*, respectively), with each video containing hundreds of particle tracks. Two types of samples, Rotor-Bridge-Tail and Bridge-Tail were tested. At least 6,000 particle tracks were analyzed for each sample and treatment condition. Only samples containing all three components, the rotor, bridge, and propeller, showed increased diffusion following ATP addition.

Within the helical propeller family, pumping was not simply present or absent, but quantitatively tunable by architecture (**Fig. 5B**, **Fig. S7A–J**). Because the helical propeller variants were adjusted to comparable axial lengths, the differences in downstream axial flux mainly reflect differences in programmed helical architecture rather than total axial extent. The 90° helical propeller variants produced the strongest rotation-pumping coupling, with opposite-handed designs driving axial flux in opposite directions, whereas the 180° and 270° variants generated substantially weaker pumping. Thus, handedness sets the direction of pumping, whereas programmed helical architecture tunes its magnitude.

To identify a general organizing principle within the helical propeller series, we projected the architectures onto an effective helical slope coordinate, P/(πD), where P is pitch and D is characteristic diameter (**Fig. 5C**). Across the helical propellers, downstream pumping increased with effective helical slope, qualitatively consistent with the trend expected for low-Reynolds-number helical pumping from resistive-force theory (*29*).

We next sought to construct a candidate helical swimmer by coupling the hh2-90-4 helical propeller module to the rotary motor PS3 F₁-ATPase. To this end, we repurposed one building block of the helical propeller as a bridge protein, retaining its N-terminal propeller-forming interface while engineering its opposite end to bind the F₁-ATPase rotor, thereby enabling attachment of the rotor to the propeller **(Fig. 5D)**. Rotor-binding motifs targeting the γ subunit were generated using BindCraft and fused to the retained propeller-interface motif using RFdiffusion to create bridge designs **(Figs. S8 and S9)**. Following biochemical screening and pull-down enrichment, nsTEM analysis supported the formation of assemblies with the intended Rotor-Bridge-Propeller helical swimmer architecture. For the assembly mediated by bridge variant C, 2D class averages and a 3D reconstruction supported the designed overall organization, with the helical propeller positioned relative to the rotor as intended **(Fig. 5E)**.

We then used single-particle tracking to test whether these candidate helical swimmer assemblies exhibited ATP-dependent changes in apparent diffusion (**Fig. 5F**). Mutated helical propeller modules containing a single cysteine were labeled with ATTO643, assembled with bridge and rotor, and imaged near a glass coverslip in PEG solution. For each sample, at least three videos were recorded from the same well before and after addition of 2 mM ATP. Across seven independent experiments with the complete Rotor-Bridge-Propeller assembly and four independent control experiments with the Bridge-Propeller sample, at least 6,000 particle tracks were analyzed for each sample and treatment condition. Apparent diffusion coefficients were estimated for each experiment by mean-squared-displacement analysis (**Fig. 5F and S10**). The complete Rotor-Bridge-Propeller sample showed an increase in mean apparent diffusion following ATP addition (**Fig. 5G**). In contrast, the Bridge-Propeller control, which lacked the rotor, showed a relative decrease over the same time interval, possibly reflecting time-dependent surface adsorption or aggregation during the single-particle tracking assay. Across replicate measurements, a positive ATP-dependent shift was observed only when all three components (propeller, bridge, and rotor) were present (**Fig. 5H**). Although this assay does not directly measure propulsion force or establish unidirectional swimming, the selective ATP-dependent increase in apparent diffusion is consistent with mechanical coupling between rotor activity and the designed helical propeller. These results support the idea that designed protein shape can be integrated with an ATP-consuming rotor to produce a new, measurable functional response.

Together, these simulations and tracking experiments define how shape relates to function for designed protein assemblies. While symmetric architectures occupy a low-pumping regime, helical propellers provide a tunable route for directional fluid transport. The rotor-coupled tracking experiments further suggest that designed protein architectures can be integrated as functional modules in ATP-responsive nanoscale assemblies. More broadly, these results establish blueprint-guided shape control as a strategy for linking programmed protein architecture to nanoscale fluid transport and provide a framework for converting designed protein shapes into functional modules of protein-based nanomachines.

## Discussion

In summary, our integration of CAD-based blueprints with diffusion-based backbone design enables programmable control over protein shape across multiple conceptual and organizational scales, spanning single-chain geometry, accessible shape space, higher-order assembly architecture, and shape-guided function. We demonstrate the generality of this framework through experimental validation and structural characterization.

Our approach of treating shape directly as an explicit design parameter is distinguished from strategies in which global shape emerges indirectly from fold topology or symmetry. This distinction is notable at the assembly level: In our approach, tunable local shape states of shared building blocks serve as architectural parameters that propagate into distinct higher-order organizations. In this fashion, the geometry of each building block can be matched with the target assembly architecture, so that local curvature and twist determine the global form of the assembly (**Fig. 4**). This continuity between building-block shape and higher-order architecture is not readily achieved by stacking predefined units or positioning components in three-dimensional space, where component shape and assembly architecture are not explicitly co-designed.

Co-design approaches are especially important when shape-encoded emergent functions arise from specific parameters of the overall architecture, for example hydrodynamic properties and motility (**Fig. 1C** panel 4). Our approach makes such complex features highly programmable and tunable at the building-block stage (**Fig. 4**). The rotor-coupled helical assembly (**Fig. 5**) provides an initial experimental demonstration of this principle.

Our framework also establishes direct interoperability between protein design and existing standard CAD representations and industrial design software. This interoperability expands the accessible protein shape space beyond architectures commonly sampled in natural protein evolution or conventional protein design, including forms inspired by biological materials. More conceptually, the ability to translate externally designed shape blueprints into protein architecture suggests a parallel between protein engineering and industrial machine design, in which function arises from programmable shape organization across scales.

Future work concerns integrating current shape-guided design with simultaneous control over residue-level local protein features, and extending the accessible size and complexity of designed assemblies. Incorporating higher-resolution constraints, multi-state shape specifications, and environmentally responsive design will further extend this framework from static architectural programming toward dynamic protein systems with controllable functions (*1*, *2*). It will be powerful to integrate the shape programmability of our framework with exciting recent approaches to design switchable assemblies (*30*), protein mechanical components (*19*), and protein walkers (*31*, *32*), as well as concurrently developed virtual-reality-based design methods (*33*). Our work provides a foundation for building a vast universe of such emergent and new-to-nature nanoscale protein functions.

## Acknowledgments

We thank the UCSF Cryo-EM Core for access to electron microscopy instrumentation and technical support; the UCSF Center for Advanced Light Microscopy for access to the Nikon ECLIPSE Ti2 inverted microscope used for single-particle tracking; and the UCSF Macromolecular Structure Group Crystallography Facility for crystallization screening and optimization. We thank Prof. Dr. Christoph von Ballmoos for kindly providing the ATP synthase-deficient Escherichia coli strain DK8.

## Author contributions

Y.Q. and T.K. developed the conceptual approach for programmable protein shape design. Y.Q. developed the computational framework and designed and screened all proteins and assemblies. Y.Q. led and performed the experimental characterization, including protein production and biochemical characterization, CD spectroscopy, negative-stain and cryo-electron microscopy, and rotor-coupled single-molecule tracking experiments, and analyzed the resulting data. G.Z. contributed to the tracking-data analysis, hydrodynamic simulations, and interpretation of the results. K.Y. contributed to the design and pilot testing of the single-molecule tracking experiments. S.L. and A.L. processed the crystallographic data and determined and refined the X-ray structures with guidance and supervision from J.S.F. K.I. and G.C. contributed to protein shape design strategies. J.S.F. provided guidance and resources for the crystallographic studies. B.H. provided guidance and resources for the single-molecule studies. T.K. provided guidance, supervision, and resources. Y.Q. wrote the manuscript, and Y.Q. and T.K. revised it with contributions from the other authors. All authors read and commented on the manuscript.

## Data, code, and materials availability

The cryo-EM density maps have been deposited in the Electron Microscopy Data Bank under accession codes EMD-78070, EMD-78072, EMD-77949, and EMD-77950. The negative-stain electron microscopy reconstructions have been deposited under accession codes EMD-77955, EMD-77954, EMD-77953, EMD-77951, EMD-77952, and EMD-77927. Code and scripts used in this study are available at https://github.com/yilunqi/Shape-conditioned-protein-design.

---

**Fig. S1.**
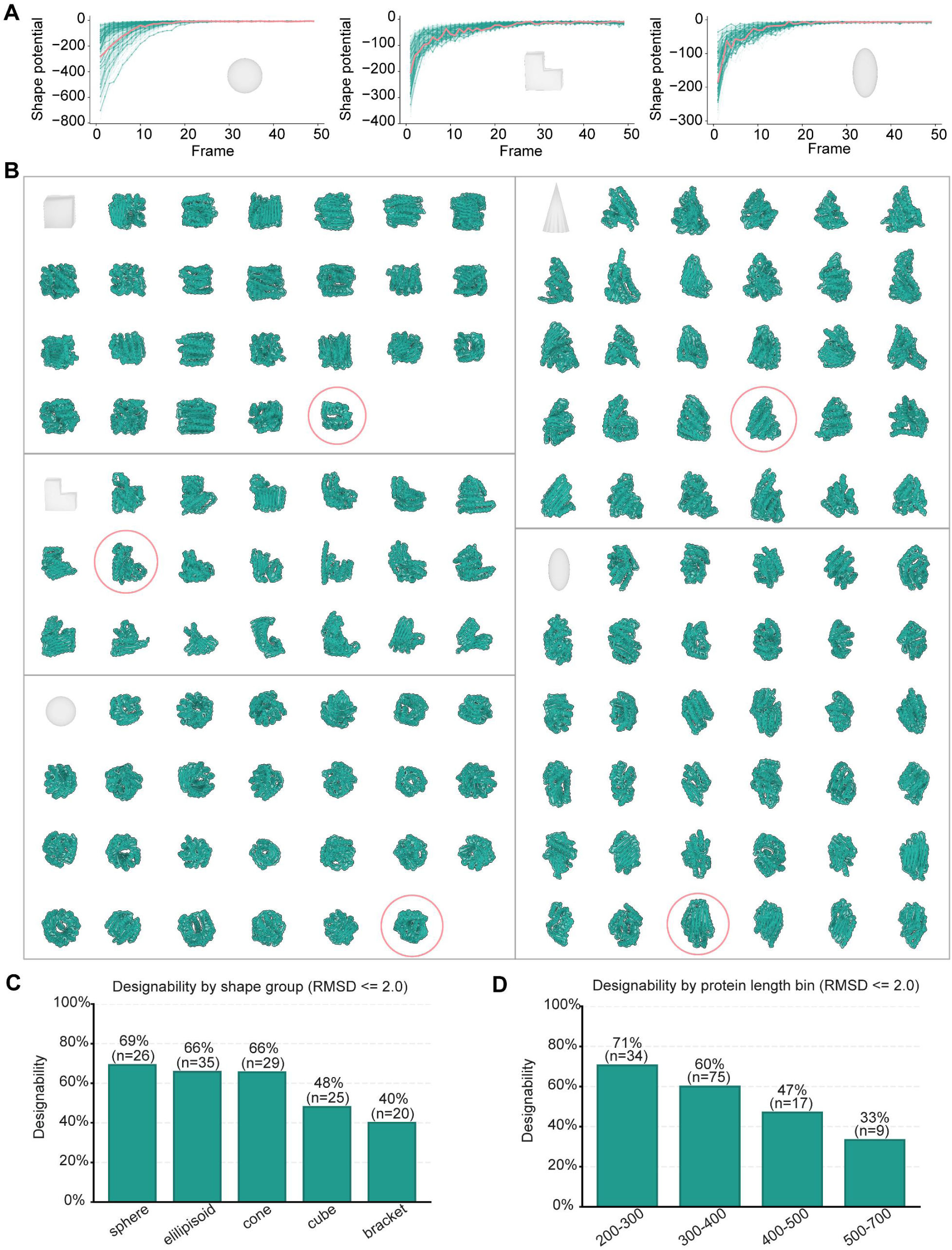
*In silico* evaluation of *de novo* shape-conditioned backbone generation and designability. (A) Shape-potential trajectories during diffusion for representative shape classes: sphere, bracket, and ellipsoid. Individual trajectories are shown in green, and designs selected for experimental testing are highlighted in orange. For each shape class, 100–200 trajectories were sampled. (B) Representative backbones matching the target shape envelopes across five blueprints: cube, cone, bracket, ellipsoid, and sphere. Backbone selection rates were 25/200 for cube, 29/200 for cone, 20/200 for bracket, 35/100 for ellipsoid, and 26/100 for sphere. (C) Designability across shape classes, defined as the fraction of ProteinMPNN-designed sequences whose ESMFold-predicted structures matched the designed backbones with Cα RMSD ≤ 2.0 Å. Eight sequences were generated per backbone using ProteinMPNN. (D) Designability across protein length, showing a decrease in *in silico* success rate with increasing chain length.

**Fig. S2.**
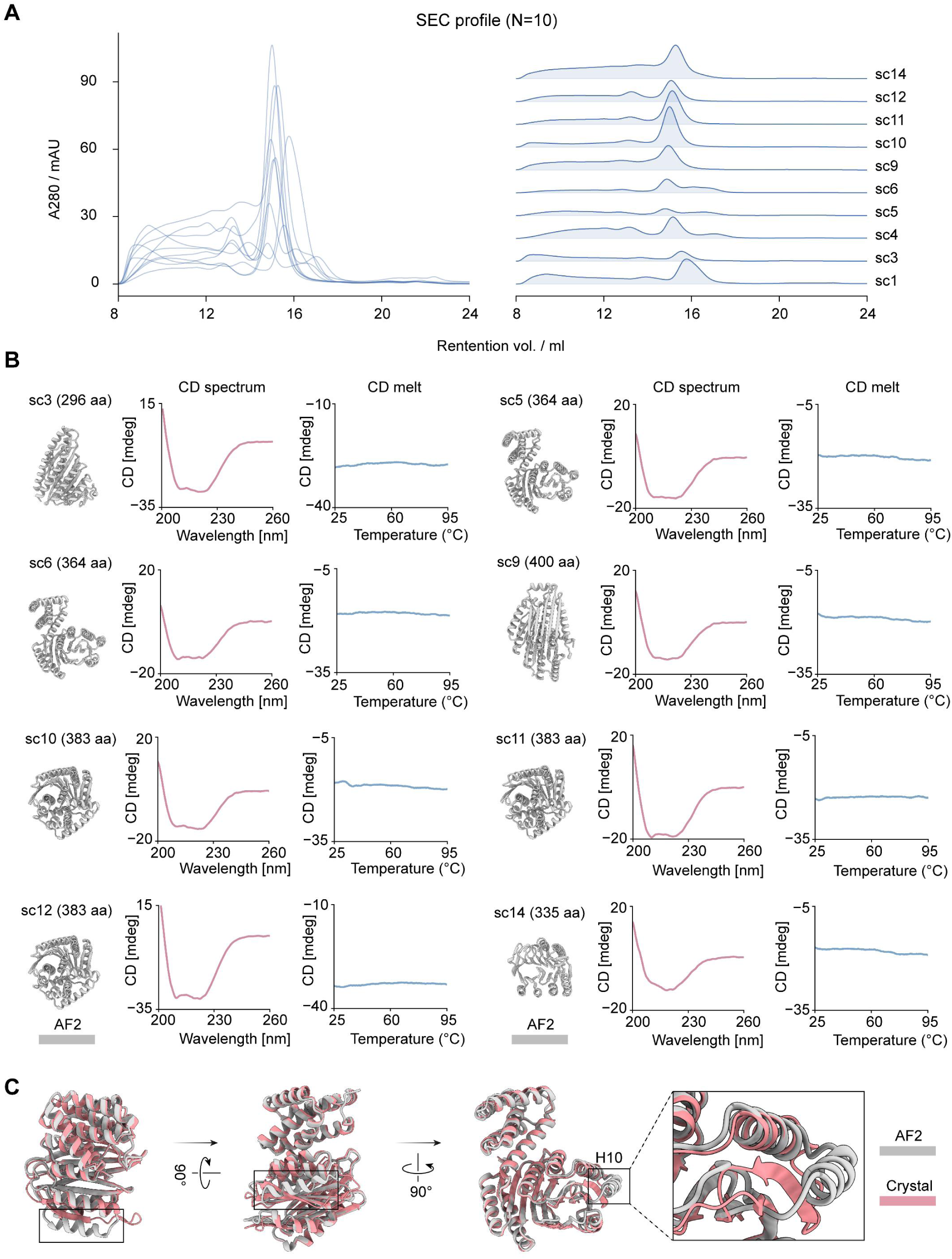
Experimental validation confirms the accuracy of *de novo* shape-conditioned designs. (A) Size-exclusion chromatography, SEC, profiles of selected designs, N = 10, showing predominantly monodisperse peaks consistent with soluble, well-behaved proteins. Three sequences were tested for each selected backbone; 10 of 15 designs were solubly expressed and analyzed by SEC. (B) CD spectra and thermal denaturation profiles (CD melts) of representative designs, indicating well-folded and thermally stable α/β structures. Corresponding AF2 models and sequence lengths are shown. (C) Structural comparison between the AF2 model (gray) and the crystal structure (pink) of the bracket design SC4. The comparison highlights a local rearrangement in which a helical segment at the edge of the design adopts a β-strand conformation in the crystal structure.

**Fig. S3.**
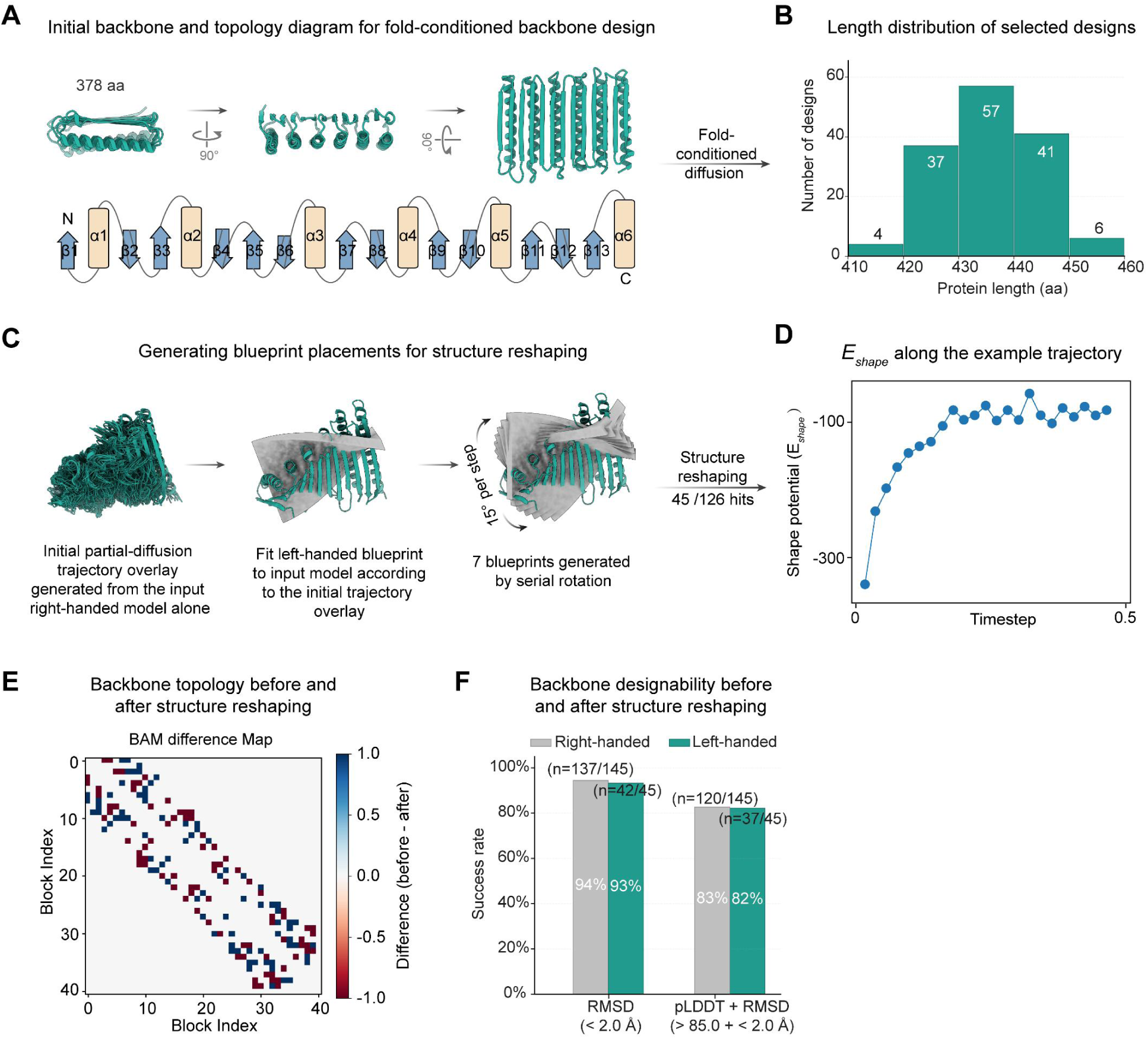
*In silico* details of the structure reshaping workflow and designability analysis. (A) Initial α/β backbone model containing a β-sheet layer, manually constructed as input for fold-conditioned RFdiffusion. (B) Length distribution of selected fold-conditioned backbones, n = 145, spanning 410–460 residues. Fold-conditioned diffusion yielded 145 of 200 backbones with the intended β-sheet layer, all of which adopted a right-handed twist. (C) Blueprint generation for structure reshaping. CAD blueprints were aligned to the noised backbone, and multiple orientations were generated by rotational sampling. For each blueprint–backbone pair, two 25-timestep trajectories were sampled, followed by sequence design using ProteinMPNN and filtering using AF2. After reshaping, 45 of 126 designs exhibited a left-handed twist. (D) Shape-potential trajectory for a representative reshaping design. (E) Topology comparison before and after reshaping. The block-adjacency difference map shows that the overall topology is preserved despite substantial changes in global shape. (F) Designability of backbones before, right-handed, and after, left-handed, reshaping. Eight sequences were generated per backbone using ProteinMPNN and evaluated by ESMFold. Success rates were 94%, right-handed, n = 137/145, and 93%, left-handed, n = 42/45, using a Cα RMSD cutoff of 2.0 Å, and 83%, right-handed, n = 120/145, and 82%, left-handed, n = 37/45, when an additional pLDDT ≥ 85 filter was applied.

**Fig. S4.**
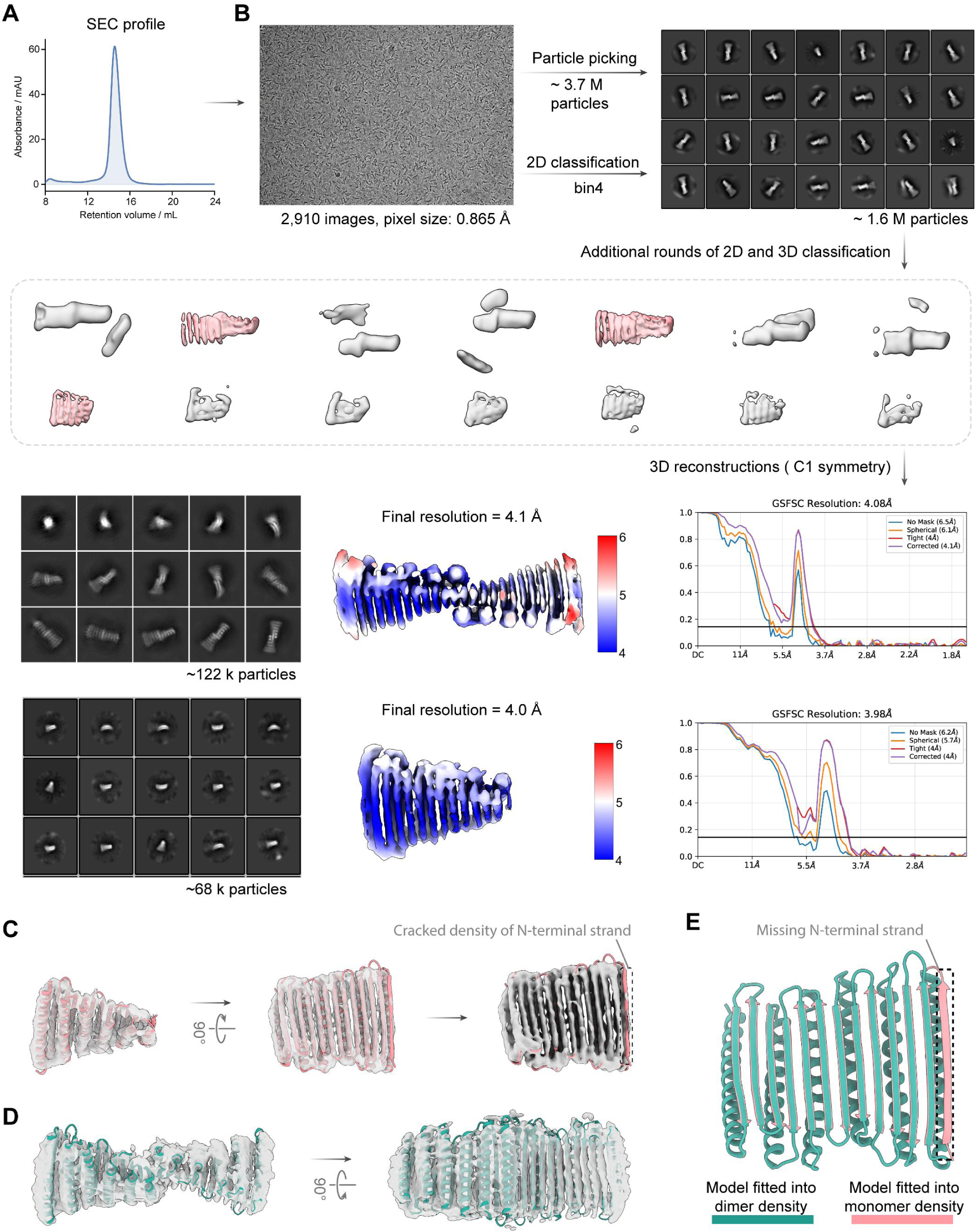
Cryo-EM validation of the left-handed twist design. (A) Size-exclusion chromatography trace of the purified sample used for cryo-EM analysis. (B) Cryo-EM data processing and reconstruction workflow. Two particle populations were refined into a 4.1 Å dimeric map, ∼122,000 particles, and a 4.0 Å monomeric map, ∼68,000 particles, both of which resolved secondary-structure features. (C) Docking of the AF2 model (pink) into the monomeric density (gray) showed overall agreement, with reduced density at the N-terminal β-strand. (D) Docking of two AF2 models into the dimeric density recapitulated the left-handed propeller architecture. (E) Comparison of the monomeric and dimeric reconstructions confirmed the left-handed twist in the dimeric state and further supported flexibility in the N-terminal β-sheet region.

**Fig. S5.**
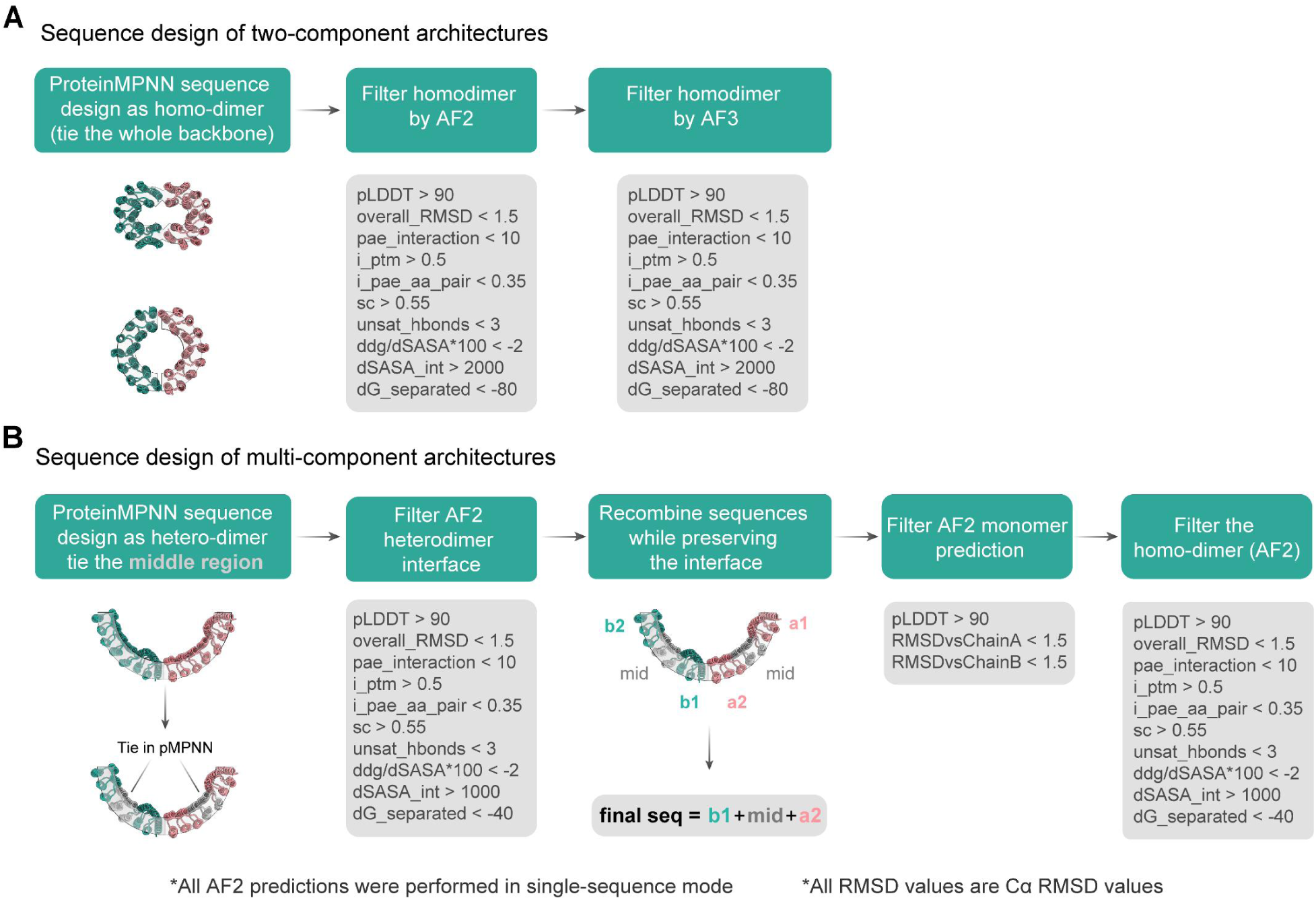
Sequence design workflows for assembly architectures. (A) Sequence design workflow for two-component dimeric assemblies. ProteinMPNN was used to design sequences for the full backbone with corresponding residues tied across the two chains. Homodimeric predictions were then filtered using AF2 and AlphaFold3 (AF3), based on model confidence, agreement with the designed backbone, and interface metrics. (B) Sequence design workflow for multi-component assemblies, with C90 shown as an example. ProteinMPNN was first applied to heterodimeric interfaces with constraints on the central region. Interface designs were filtered using AF2, and selected sequence segments were recombined to preserve the designed interfaces. Recombined sequences were then evaluated at both the monomeric and homodimeric levels using AF2. Filtering thresholds included pLDDT, Cα RMSD, and interface-confidence metrics.

**Fig. S6.**
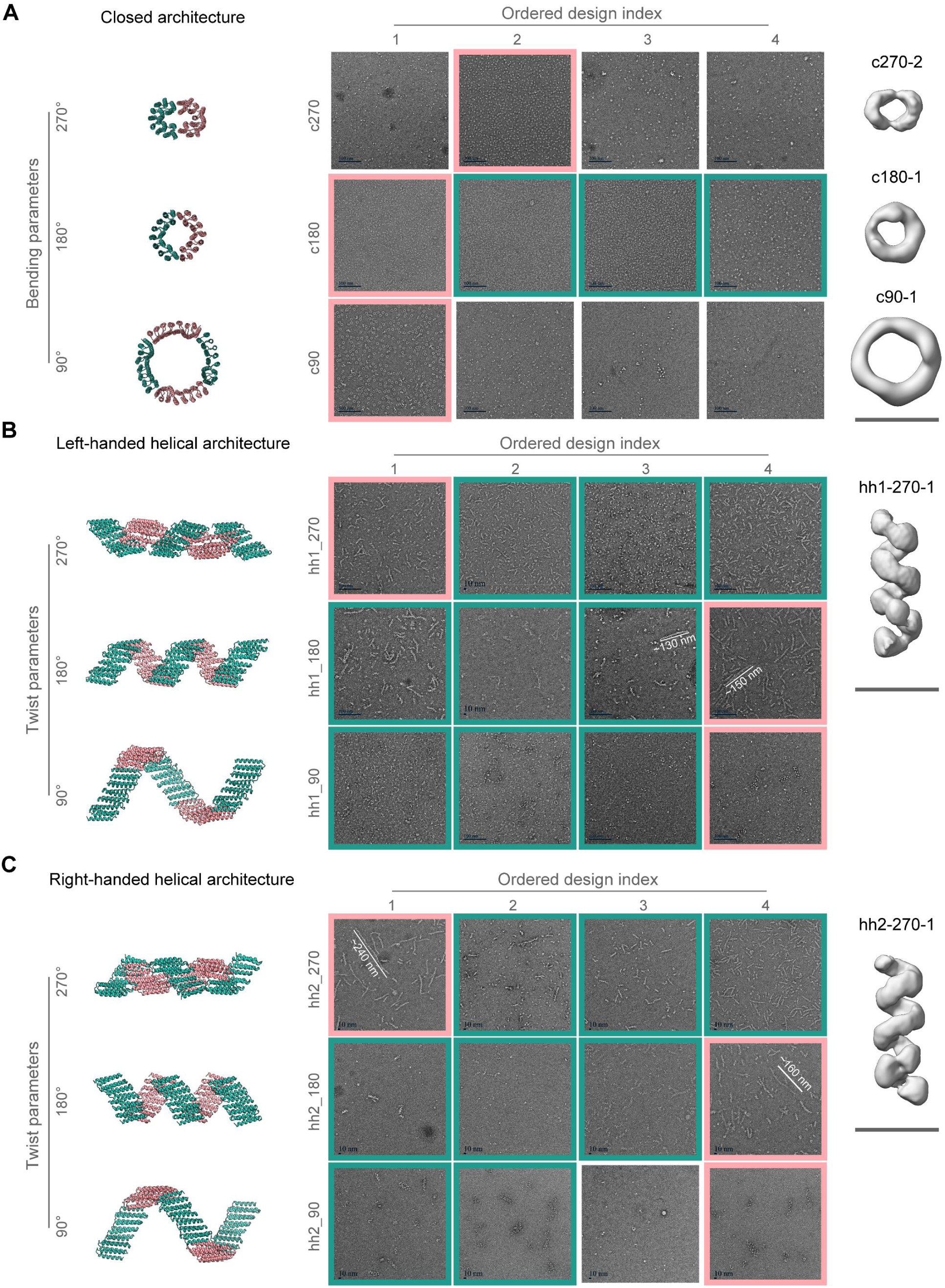
Negative-stain TEM validates the designed assembly architectures. (A-C) Experimental validation of designed assemblies by negative-stain transmission electron microscopy (TEM). Four sequences were tested for each architecture. (A) Closed assemblies with varying curvature parameters, C270, C180, and C90. (B) Left-handed helical assemblies with varying twist parameters. (C) Right-handed helical assemblies with corresponding twist parameters. Representative micrographs are shown for each design (samples were imaged directly after Ni-NTA purification). Green and pink boxes indicate designs consistent with the target architecture. Designs outlined in pink were selected for further analysis. Three-dimensional reconstructions from negative-stain TEM data, shown on the right, confirm agreement with the designed architectures. Scale bars, 10 nm.

**Fig. S7.**
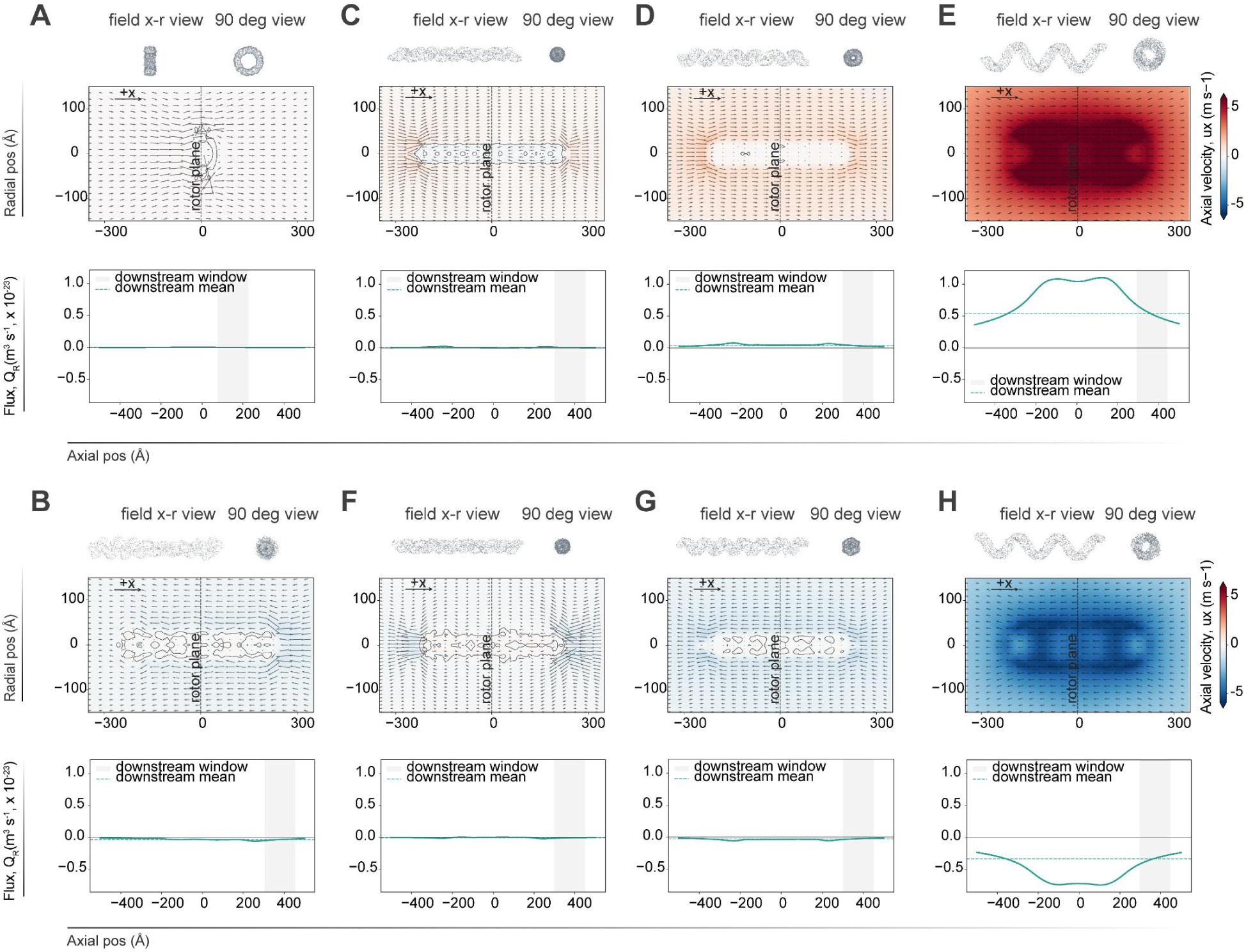
Hydrodynamic simulations across designed propeller architectures. Stokes-flow simulations of rotating protein assemblies spanning symmetric, helical, and filamentous architectures. For each structure, the upper panels show the rotating assembly in the axial-radial (***x***-***r***) flow-field view and a corresponding 90° rotated view. Colors indicate the axial velocity component, ***u_x_***, and arrows show in-plane velocity vectors in the axial-radial section. Lower panels show the corresponding finite-disk sectional axial flux profile, ***Q_R_*(*x*)/Ω**, along the rotation axis. Gray shaded regions indicate the downstream averaging window used for quantification in Fig. 5. The upper flow-field panels for (A), (B), (E), and (H) are also shown in Fig. 5A and are included here for completeness. (A) Symmetric ring control, c90. (B) F-actin control assembly. (C) Left-handed helical propeller, 270° variant, hh1-270. (D) Left-handed helical propeller, 180° variant, hh1-180. (E) Left-handed helical propeller, 90° variant, hh1-90. (F) Right-handed helical propeller, 270° variant, hh2-270. (G) Right-handed helical propeller, 180° variant, hh2-180. (H) Right-handed helical propeller, 90° variant, hh2-90. (Q_R(x)) denotes local finite-disk sectional axial flux rather than conserved flux across an infinite cross-section.

**Fig. S8.**
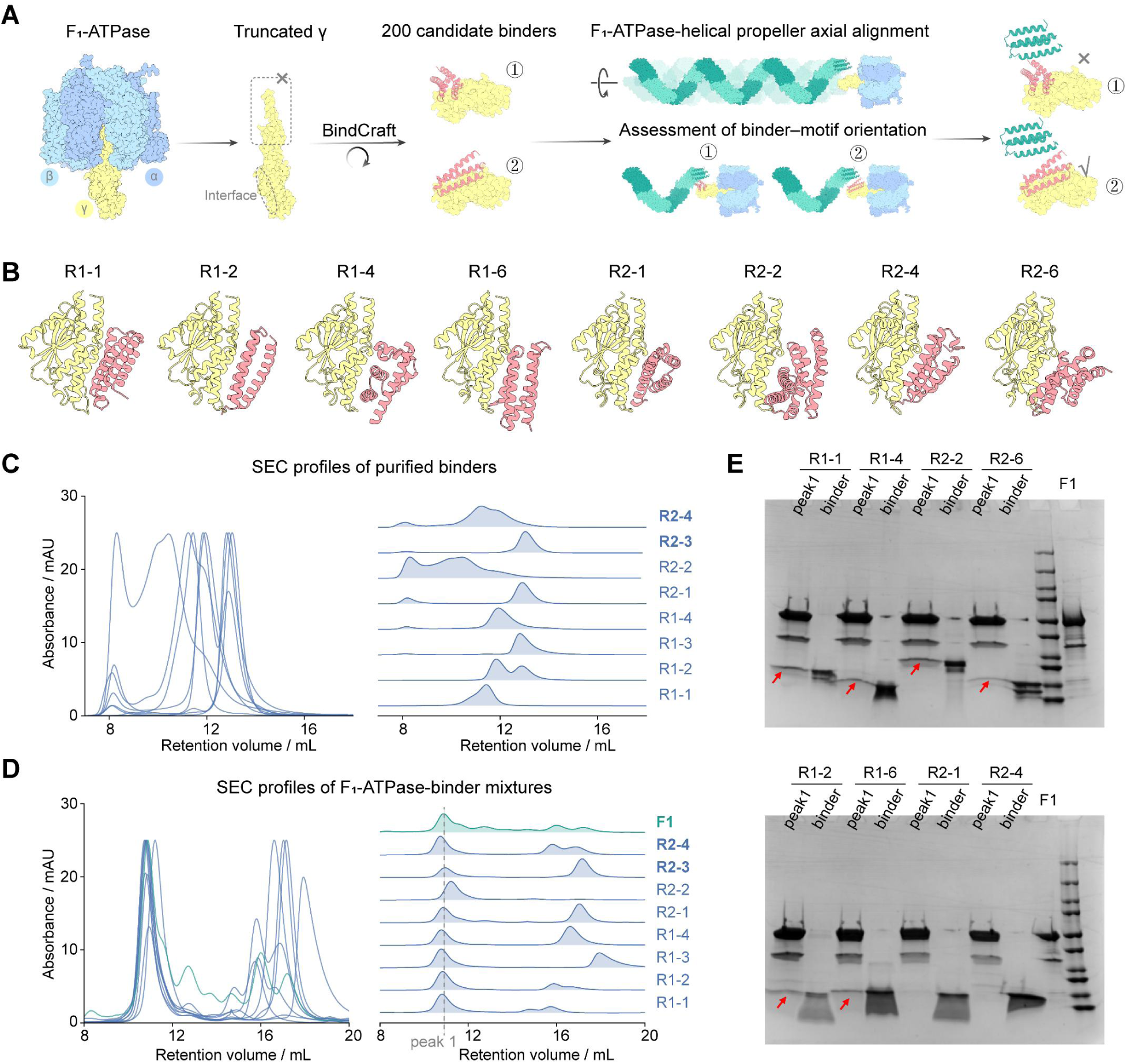
Design and validation of binders targeting the γ subunit of PS3 F₁-ATPase. (A) Design workflow for binders targeting the γ subunit of PS3 F₁-ATPase. Candidate binders generated by BindCraft were evaluated based on their orientation relative to the helical propeller after axial alignment with the F₁-ATPase model. (B) Representative binder designs generated by BindCraft from two size classes, large and small. (C) SEC profiles of purified binders, showing well-defined peaks for selected candidates. (D) SEC analysis of binder–F₁-ATPase mixtures. Purified binders were incubated with F₁-ATPase, green trace, and the mixtures were analyzed by SEC. Fractions corresponding to the F₁-ATPase peak, peak 1, were collected for further analysis. (E) SDS-PAGE analysis of SEC fractions confirming co-elution of selected binders with F₁-ATPase. Binders detected in peak 1 fractions were considered successful F₁-ATPase binders, including big1, big2, big4, big6, small2, and small6.

**Fig. S9.**
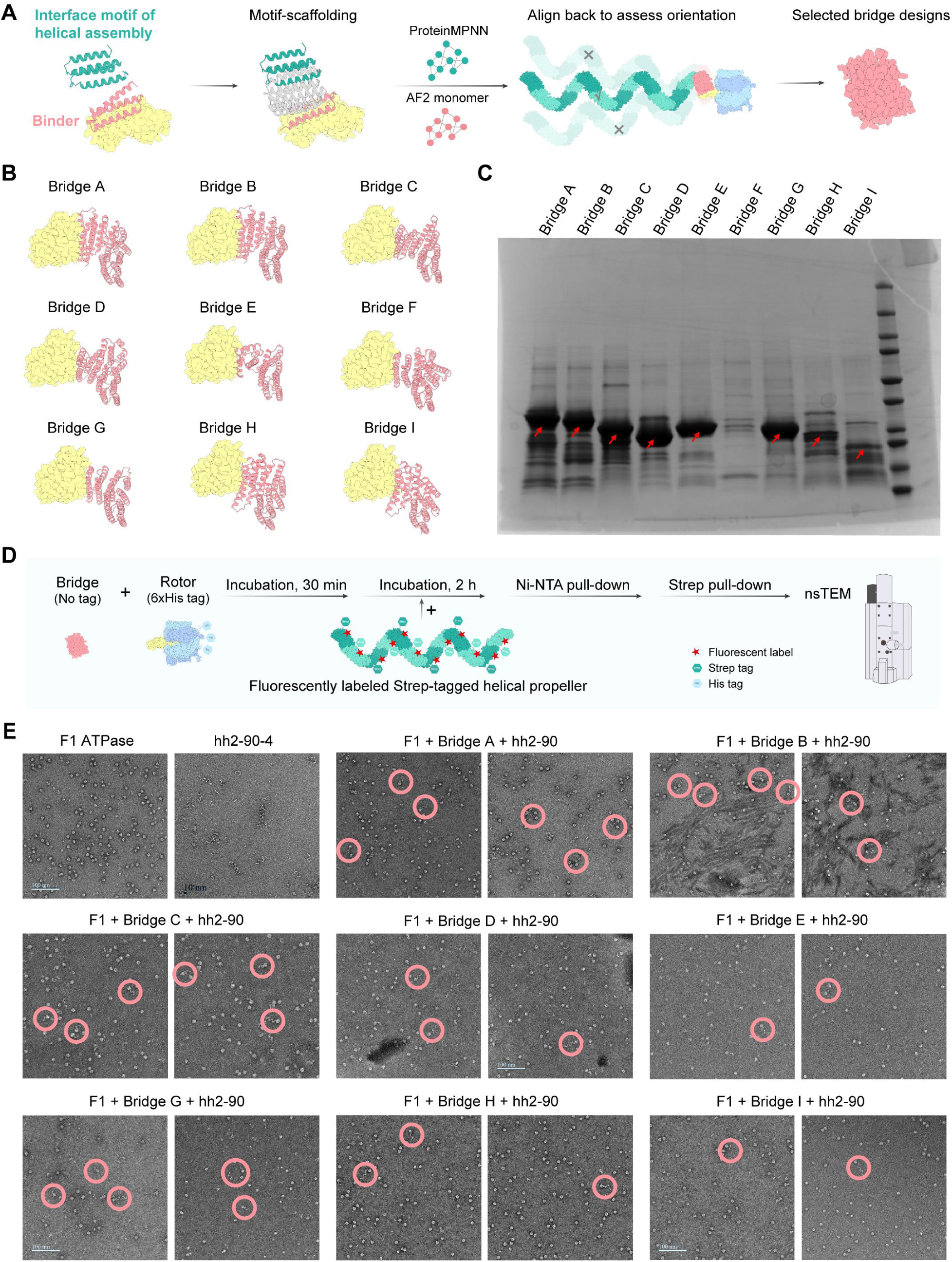
Design and validation of bridge designs for helical swimmer assembly. (A) Bridge design by motif scaffolding. Binding motifs from selected γ-subunit binders and interface motifs from the helical assembly were fused using RFdiffusion, followed by sequence design using ProteinMPNN and filtering using AF2. Designed bridges were aligned back to the full motor–propeller model to assess orientation compatibility and axial alignment between F₁-ATPase and the helical propeller. (B) Representative bridge designs, Bridge A–I, connecting the F₁-ATPase γ subunit to the helical propeller. (C) SDS-PAGE analysis of bridge expression, showing bands near the expected molecular weights for the designed bridge constructs. (D) In vitro assembly workflow. Purified F₁-ATPase, bridge protein, and Strep-tagged helical propeller were sequentially assembled and enriched by Ni-NTA pull-down followed by Strep-tag affinity purification. (For single-particle tracking experiments, a fluorescently labeled, Strep-tagged helical propeller was used.) Assembled complexes were subjected to structural characterization by nsTEM. (E) nsTEM images of assembly mixtures. F₁-ATPase and helical propeller alone are shown as references, top left. Samples containing bridge designs showed particles consistent with assembled complexes, with representative examples highlighted by red circles. Two representative micrographs are shown for each bridge design. Assemblies displaying coherent overall architecture were selected for further analysis.

**Fig. S10.**
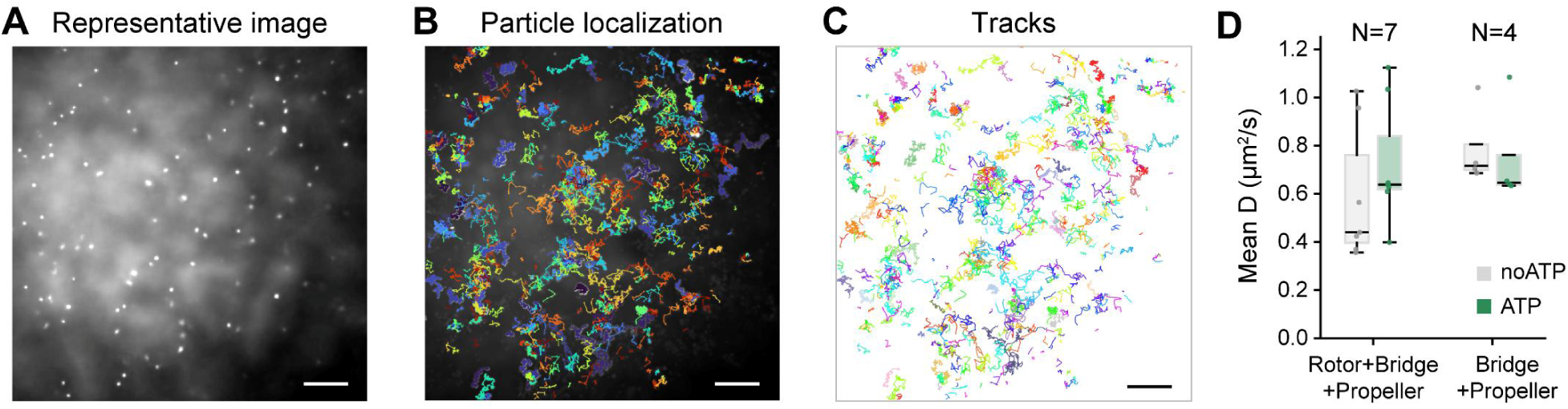
Representative single-particle tracking analysis video. (A) Representative frame from a raw video showing fluorescent particles. (B) Particle localization and trajectory overlay for the representative video. (C) Extracted single-particle tracks used for downstream diffusion analysis. Scale bars, 10 μm (A-C). (D) Mean diffusion coefficient before and after ATP addition. N denotes the number of independent experiments.

**Table S1.** X-ray data and structure statistics.

| Statistic | SC1 | SC4 |
| --- | --- | --- |
| Data collection |  |  |
| Beamline | SSRL BL12-1 | ALS 8.3.1 |
| Wavelength (Å) | 0.97946 | 1.11583 |
| Resolution range (Å) | 63.96-2.30 (2.40-2.30) | 117.31-1.68 (1.71-1.68) |
| Space group | P212121 | P212121 |
| Unit-cell parameters (Å, °) | a = 60.99, b = 142.63, c = 143.12; $\alpha = \beta = \gamma = 90$ | a = 40.98, b = 69.11, c = 117.31; $\alpha = \beta = \gamma = 90$ |
| Total observations | 561,399 (18,842) | 458,831 (11,536) |
| Unique reflections | 41,213 (1,486) | 38,432 (1,659) |
| Multiplicity | 13.6 (12.7) | 11.9 (7.0) |
| Completeness (%) | 73.1 (21.1) spherical; 84.5 (39.1) ellipsoidal | 98.6 (84.4) |
| Mean I/$\sigma$(I) | 3.6 (0.1) | 22.6 (0.7) |
| Wilson B factor (Å<sup>2</sup>) | 56.12 | 39.40 |
| Rmerge | 0.162 (2.126) | 0.045 (1.920) |
| Rmeas | 0.169 (2.216) | 0.047 (2.072) |
| Rpim | 0.046 (0.615) | 0.014 (0.749) |
| CC1/2 | 0.994 (0.542) | 1.000 (0.422) |
| STARANISO principal diffraction limits (Å) | 2.92, 1.90, 2.23 along a*, b*, c* | - |
| Refinement |  |  |
| Refinement resolution range (Å) | 63.929-2.700 (2.797-2.700) | 30.215-1.720 (1.781-1.720) |
| Reflections used in refinement | 25,161 (809) | 34,249 (3,236) |
| Reflections used for Rfree | 930 (35) | 1,822 (179) |
| Refinement completeness (%) | 74.35 (24.46) | 99.51 (95.34) |
| Rwork | 0.2373 (0.3874) | 0.2290 (0.3555) |
| Rfree | 0.3057 (0.3887) | 0.2700 (0.3795) |
| Model composition |  |  |
| Protein molecules per asymmetric unit | 4 | 1 |
| Protein residues | 1176 | 355 |
| Non-hydrogen atoms | 8845 | 2762 |
| Protein atoms | 8827 | 2725 |
| Ligands / ions | 6 Cl<sup>-</sup> | 1 Cl<sup>-</sup> |
| Water molecules | 12 | 36 |
| Model geometry and validation |  |  |
| RMSD bonds (Å) | 0.00 | 0.01 |
| RMSD angles (°) | 0.64 | 0.87 |
| Ramachandran favored (%) | 92.86 | 98.57 |
| Ramachandran allowed (%) | 6.11 | 1.43 |
| Ramachandran outliers (%) | 1.03 | 0.00 |
| Rotamer outliers (%) | 12.12 | 3.24 |
| Clashscore | 9.00 | 0.71 |
| MolProbity score | 2.76 | 1.12 |
| Values in parentheses are for the highest-resolution shell. |  |  |
| SC1 data were anisotropically truncated with STARANISO using a local weighted CC1/2 threshold of 0.30. |  |  |

**Table S2.** Cryo-EM structure statistics.

|  |  |  |  |  |
| --- | --- | --- | --- | --- |
| Parameter | Left-handed twist $\alpha/\beta$ backbone | | hh1-90-4 | hh2-90-4 |
| EMDB ID | EMD-78070 | EMD-78072 | EMD-77949 | EMD-77950 |
| Sample / Condition | Dimer | Monomer | Oligomer | Oligomer |
| Microscope | Talos Arctica |  | Talos Arctica | Glacios |
| Voltage (kV) | 200 |  |  |  |
| Detector | GATAN K3 |  |  |  |
| Pixel size (Å) | 0.865 |  | 1.73 | 1.46 |
| Electron exposure (e<sup>-</sup>/Å<sup>2</sup>) | ~ 64 |  |  |  |
| Defocus range (μm) | 1.0 ~ 2.5 |  |  |  |
| Symmetry imposed | C1 |  |  |  |
| Initial particle images | ~1,600,000 |  | ~1,200,000 | 950,081 |
| Final particle images | 122,176 | 68,134 | 74,764 | 189,388 |
| Map resolution (Å) | 4.1 | 4.0 | 7.7 | 10.0 |
| FSC threshold | 0.143 |  |  |  |
| Map sharpening B-factor | -69.8 | -295.2 | -691.7 | −100 |
| Reconstruction software | RELION 5.0/cryosparc 4.5 |  | cryosparc 4.5 |  |
| Model building | Rigid-body fitting in UCSF ChimeraX |  |  |  |

**Table S3.** nsTEM structure statistics.

| Parameter | c270-2 | c180-1 | c90-1 | hh1-270-1 | hh2-270-1 | swimmer |
| --- | --- | --- | --- | --- | --- | --- |
| EMDB ID | EMD-77955 | EMD-77954 | EMD-77953 | EMD-77951 | EMD-77952 | EMD-77927 |
| Microscope | FEI T12 |  |  |  |  |  |
| Voltage (kV) | 120 |  |  |  |  |  |
| Detector | Gatan Rio 4k |  |  |  |  |  |
| Pixel size (Å) | 1.34 | 1.34 | 2.68 | 2.68 | 2.68 | 4.5 |
| Symmetry imposed | C1 |  |  |  |  |  |
| Particles used | 1,109 | 1,566 | 1,641 | 6,156 | 6,940 | 2,380 |
| Estimated resolution (Å) | 13.2 | 21.4 | 19.6 | 15.5 | 16.4 | 23 |
| Reconstruction software | cryosparc 4.5 |  |  |  |  |  |

**Table S4.**
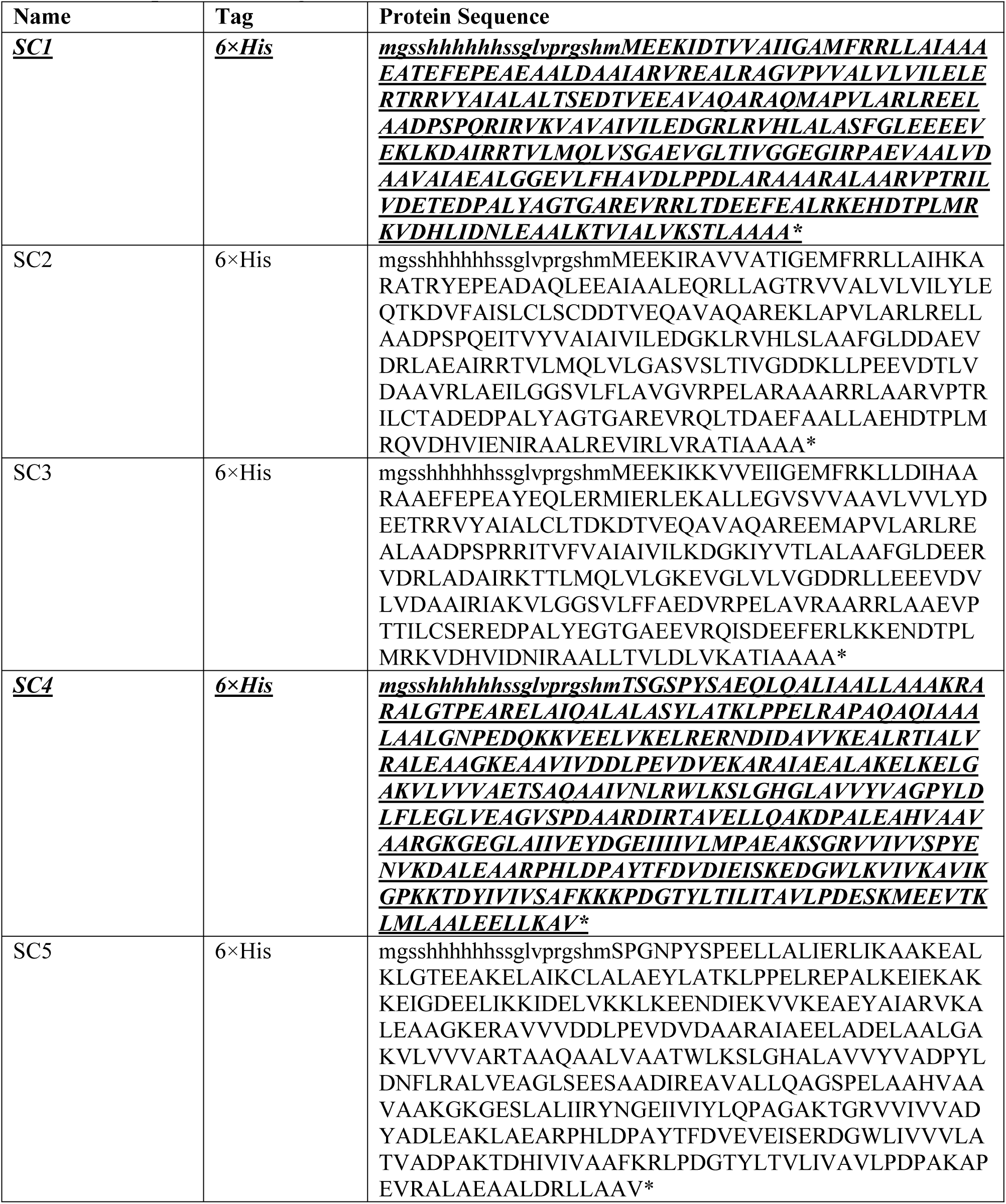
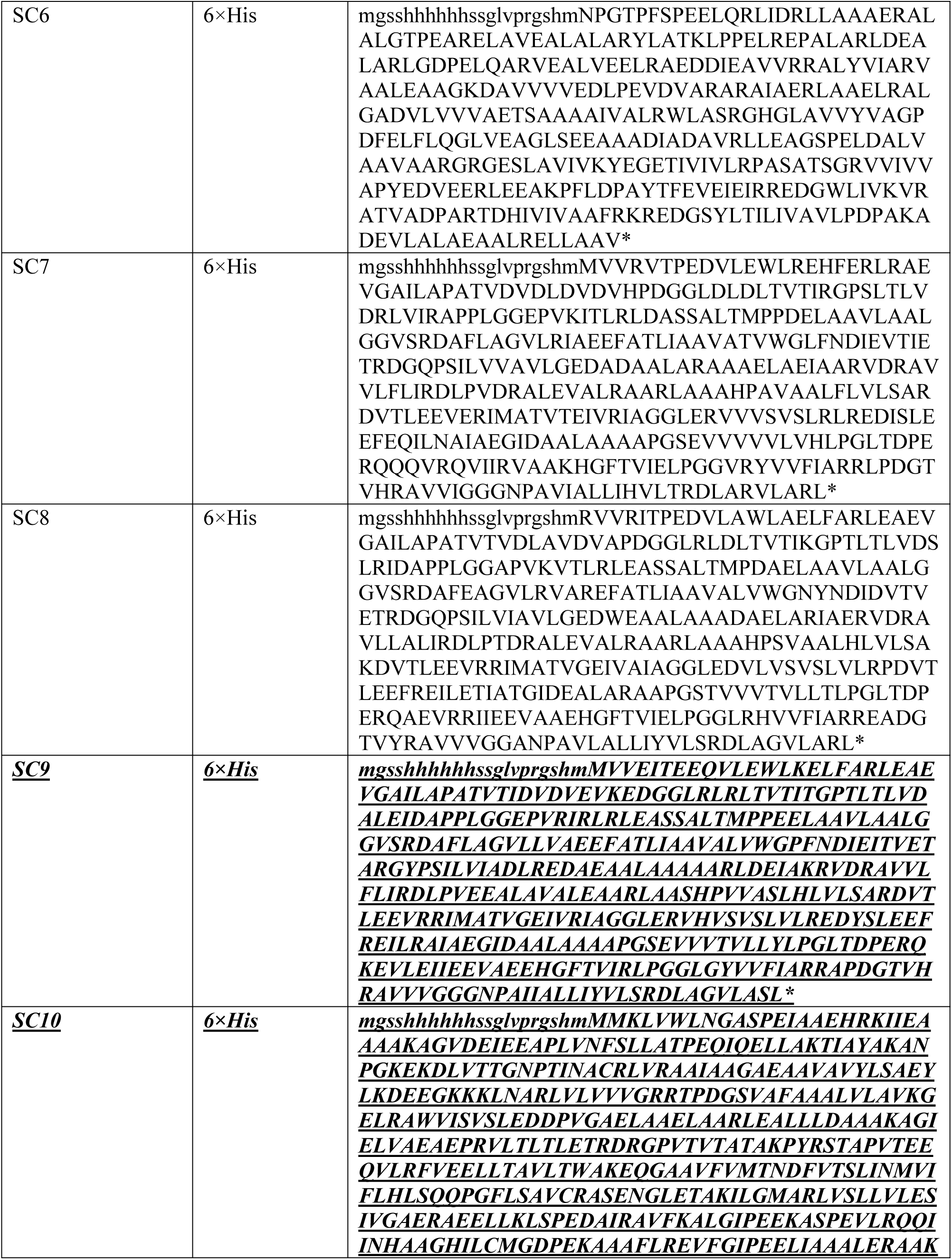
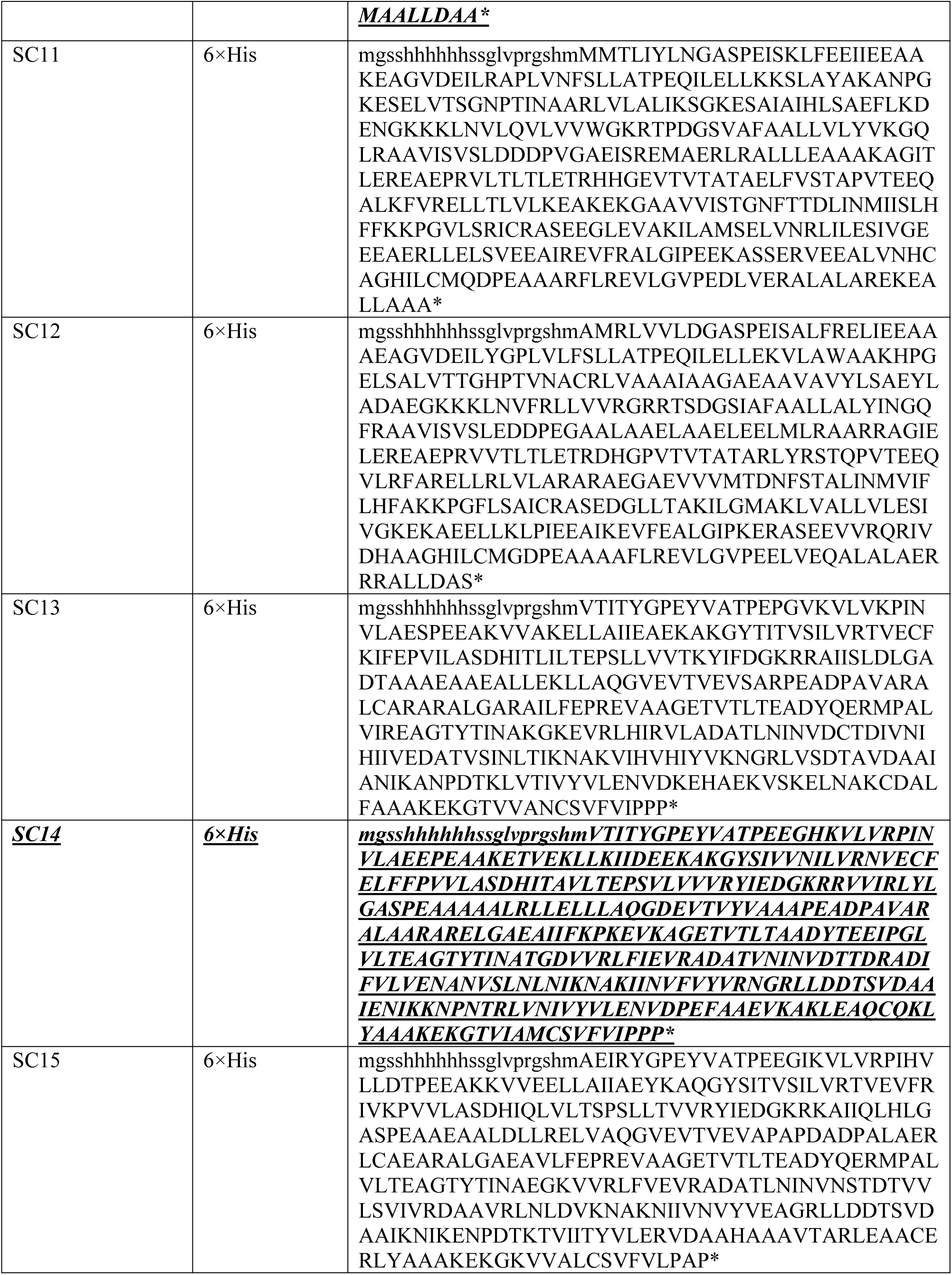
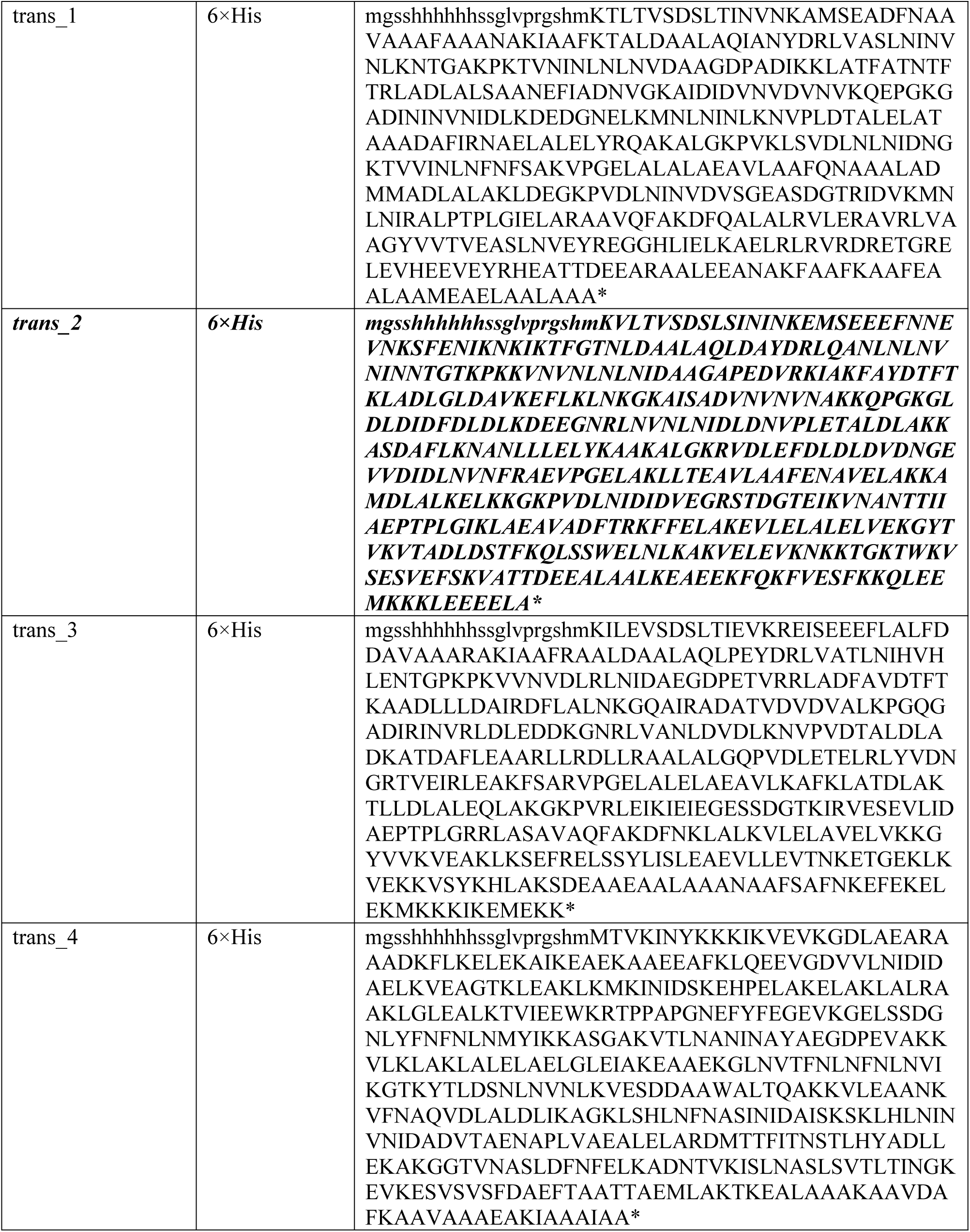
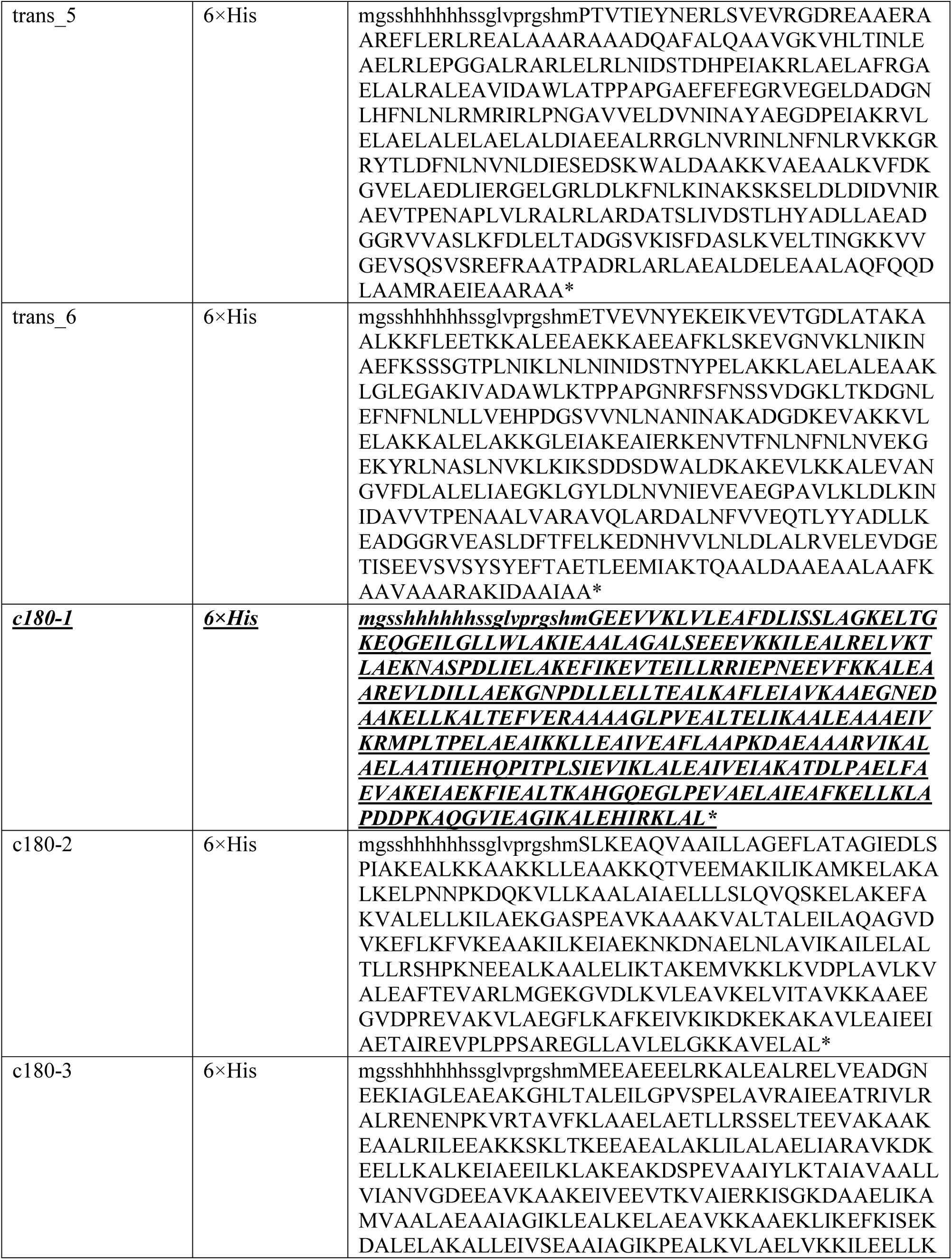
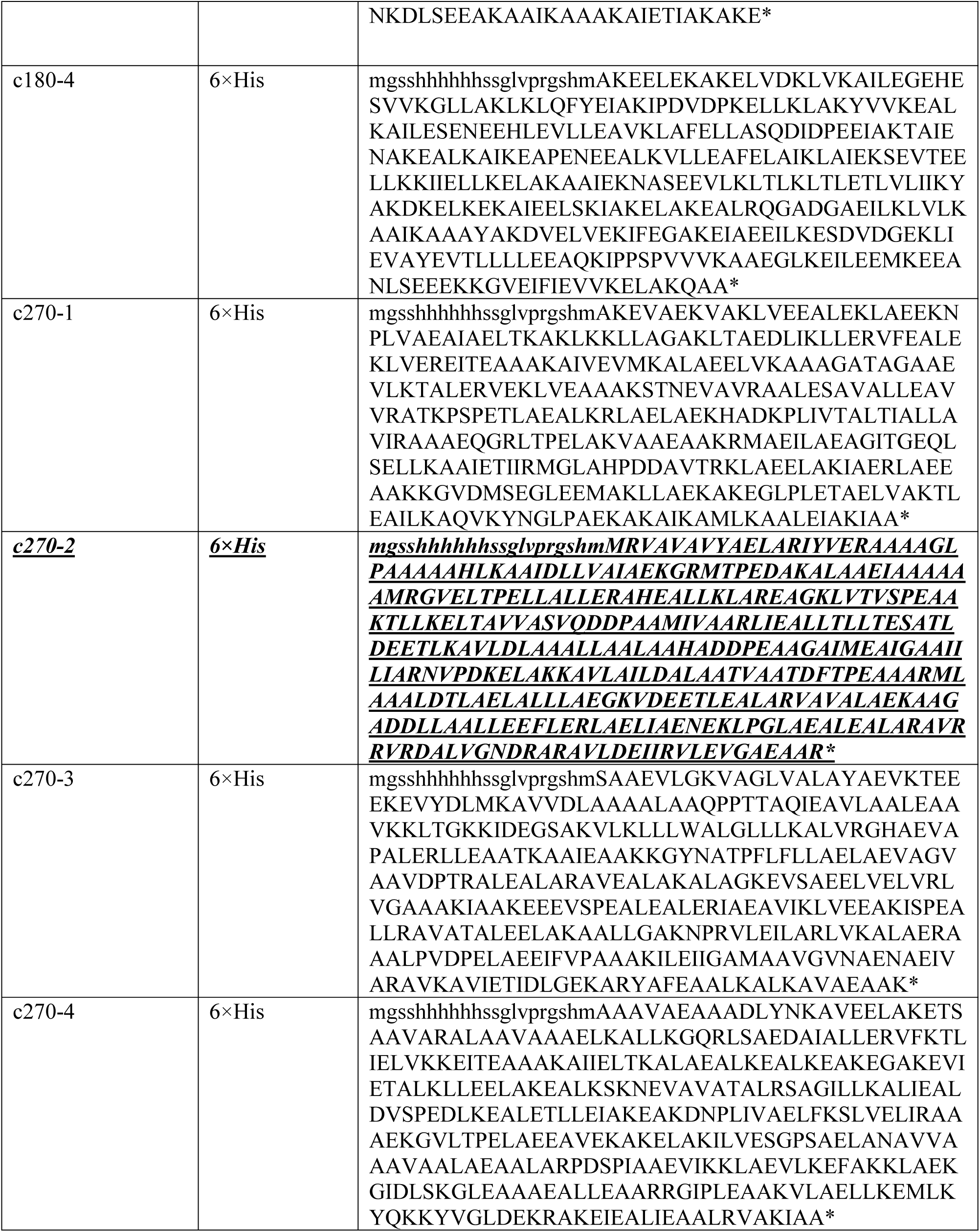
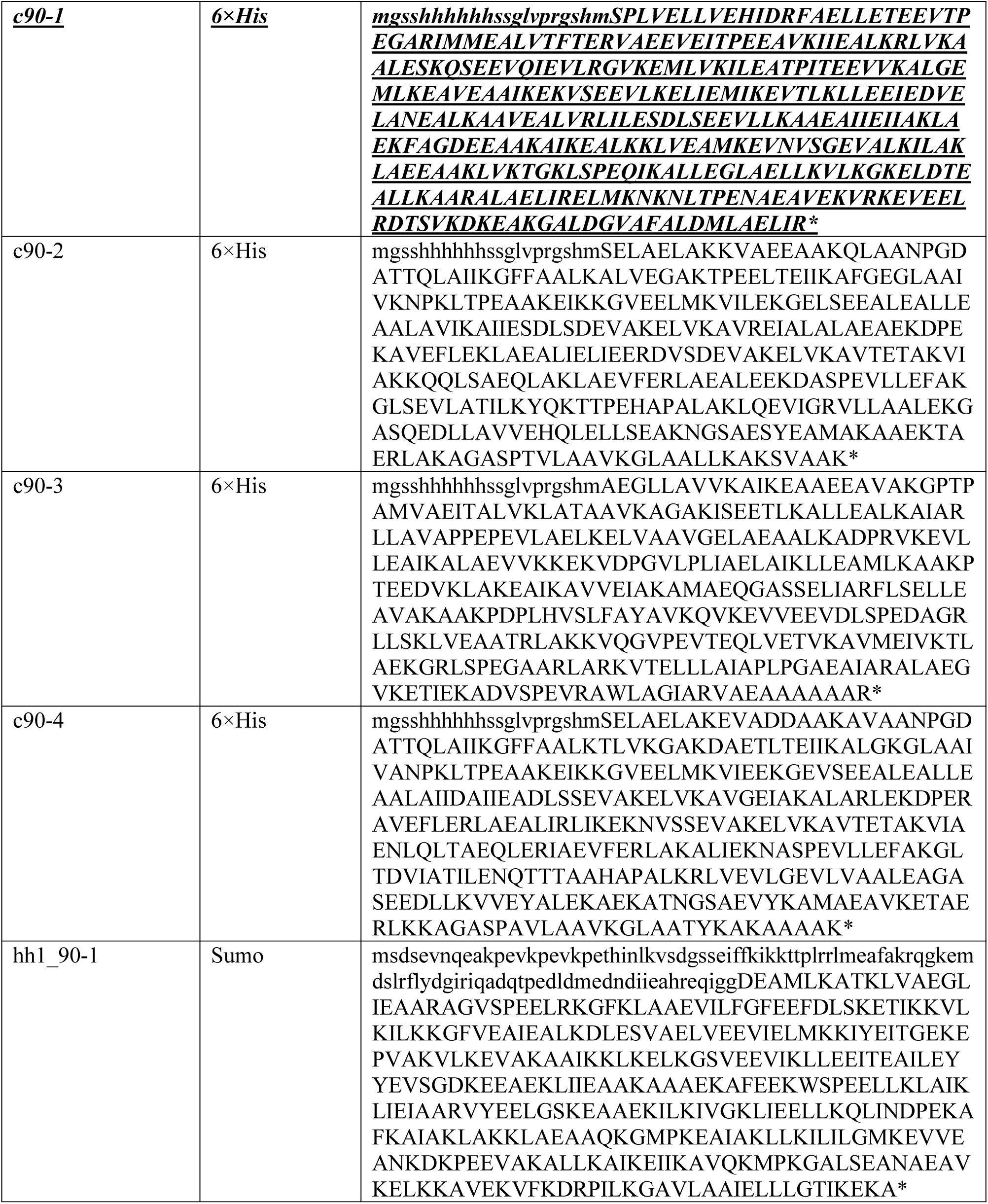
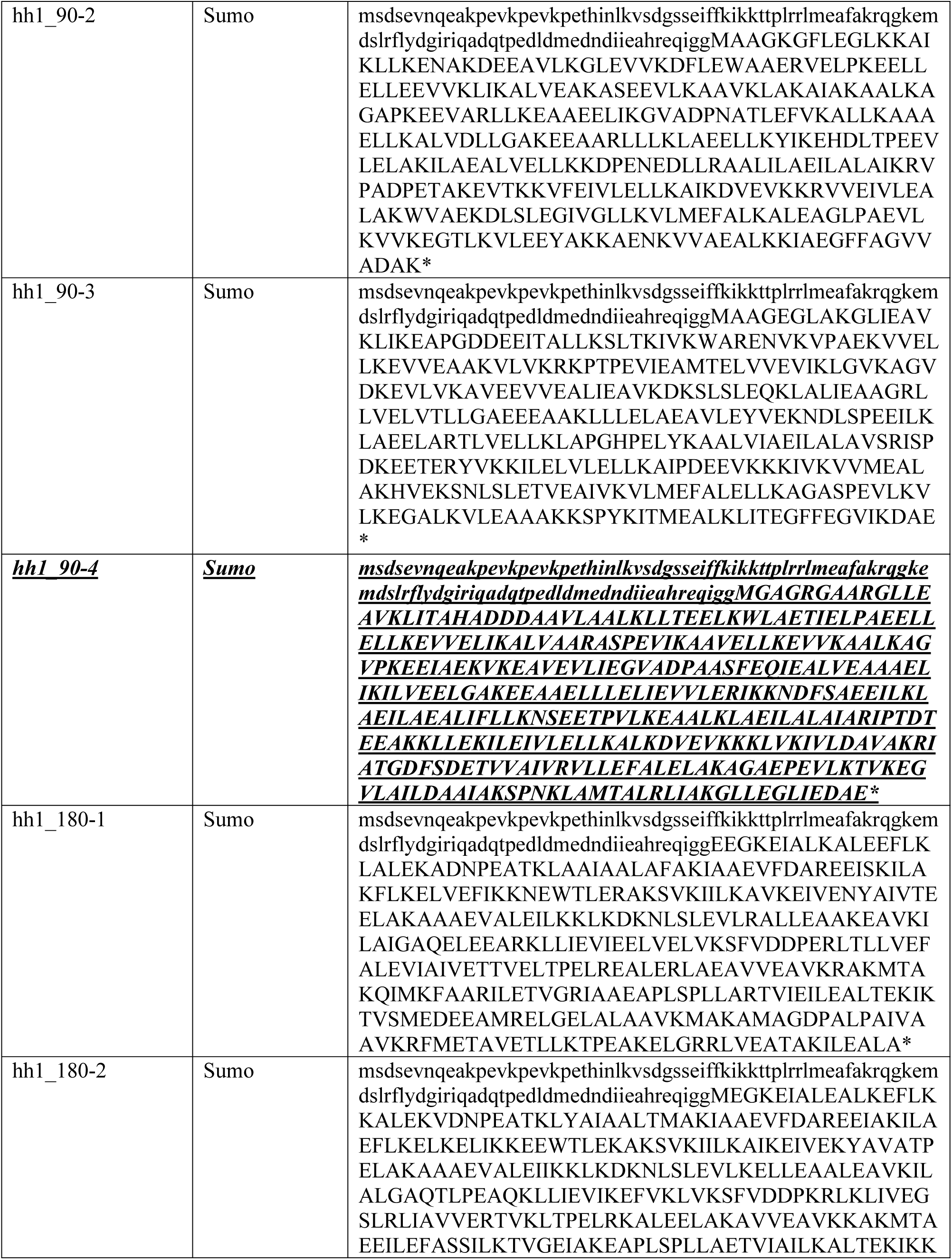
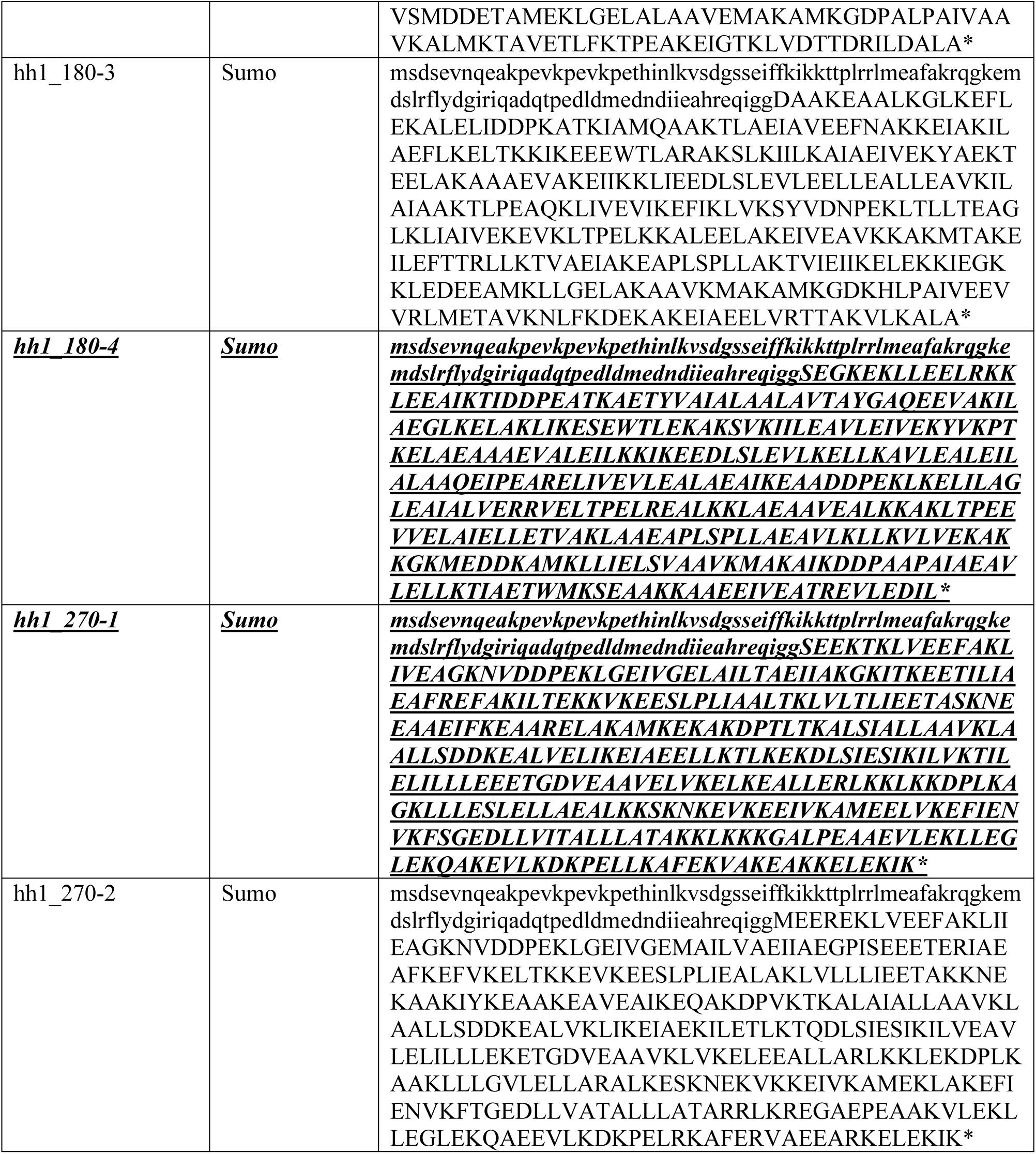
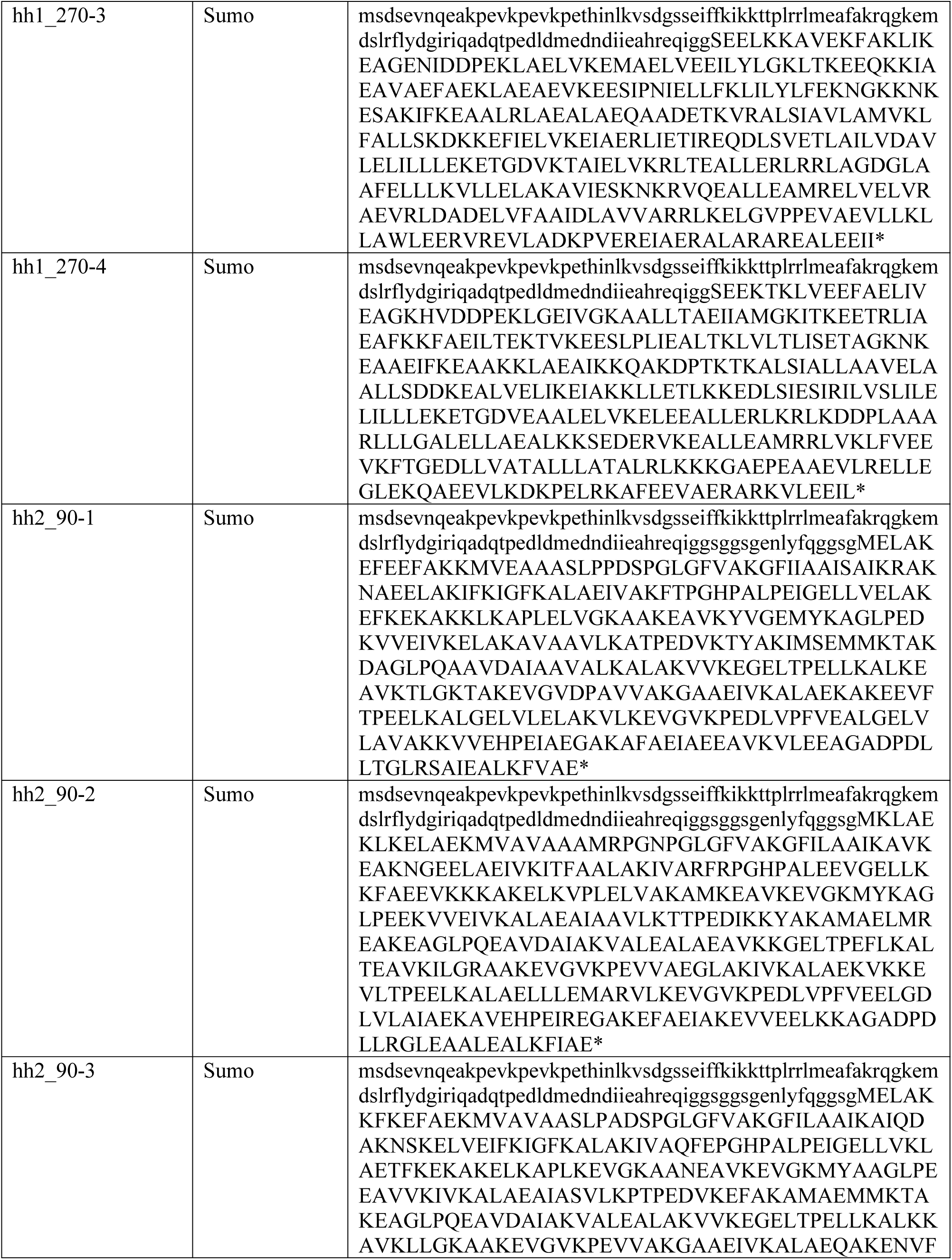
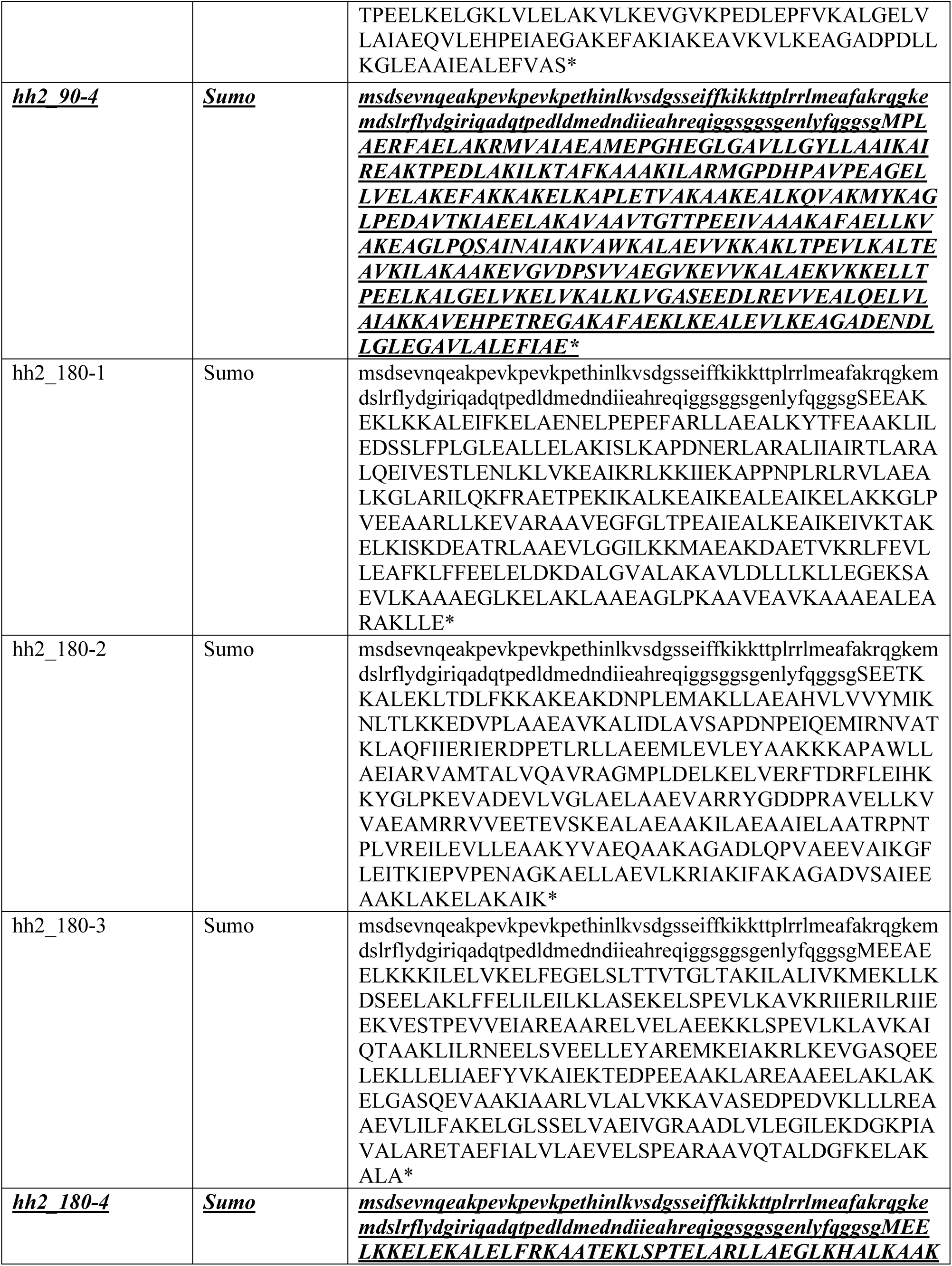
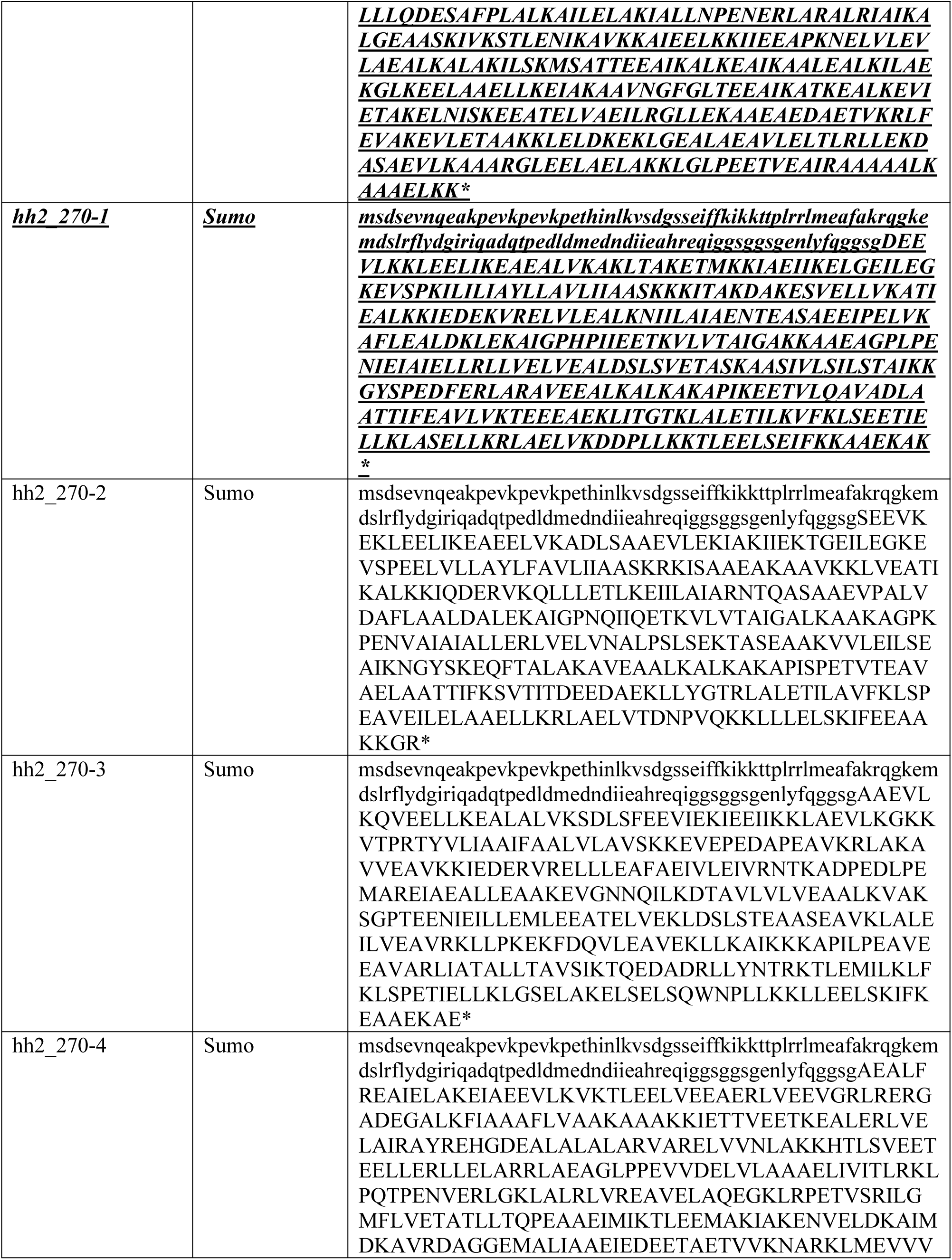
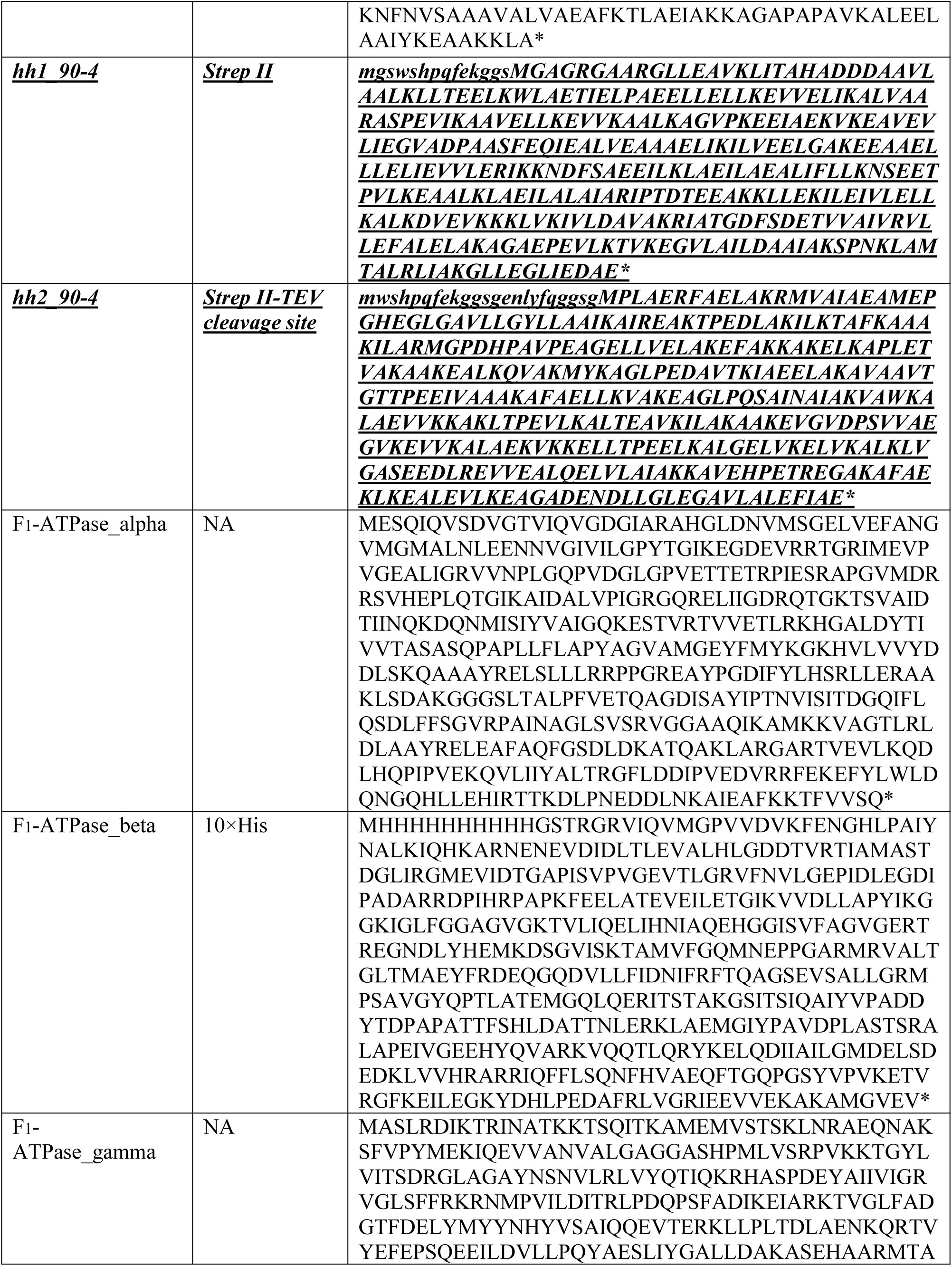
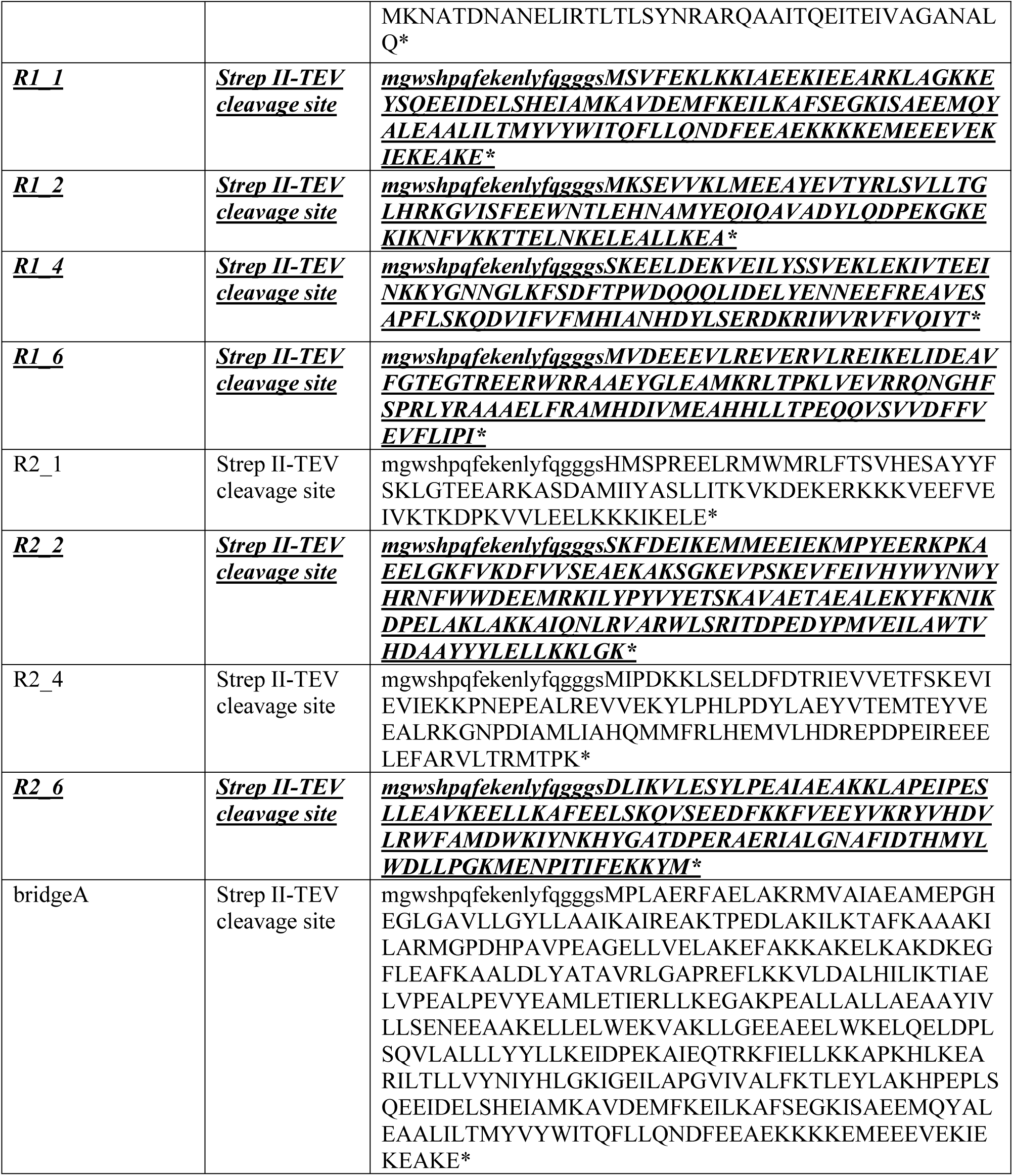
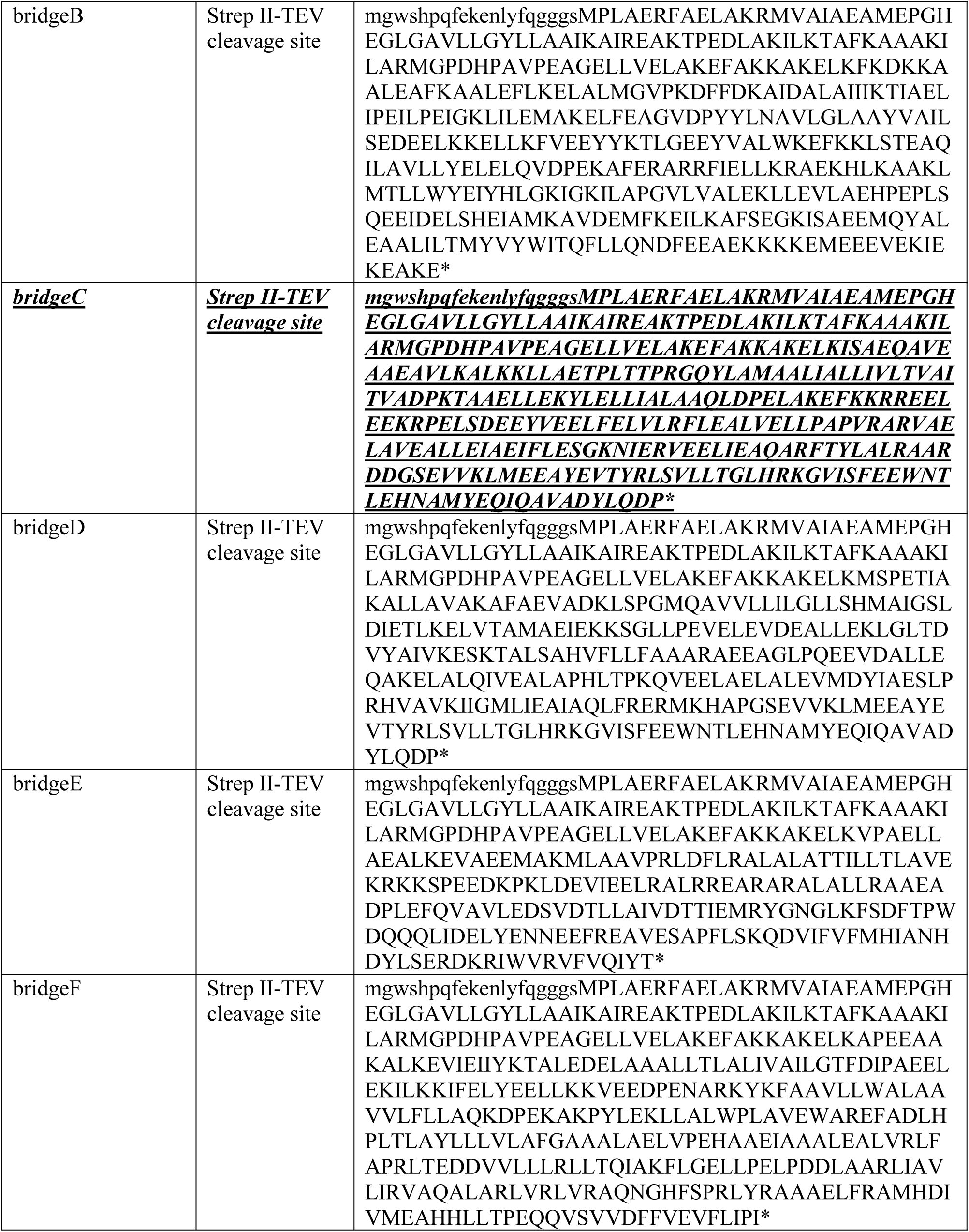
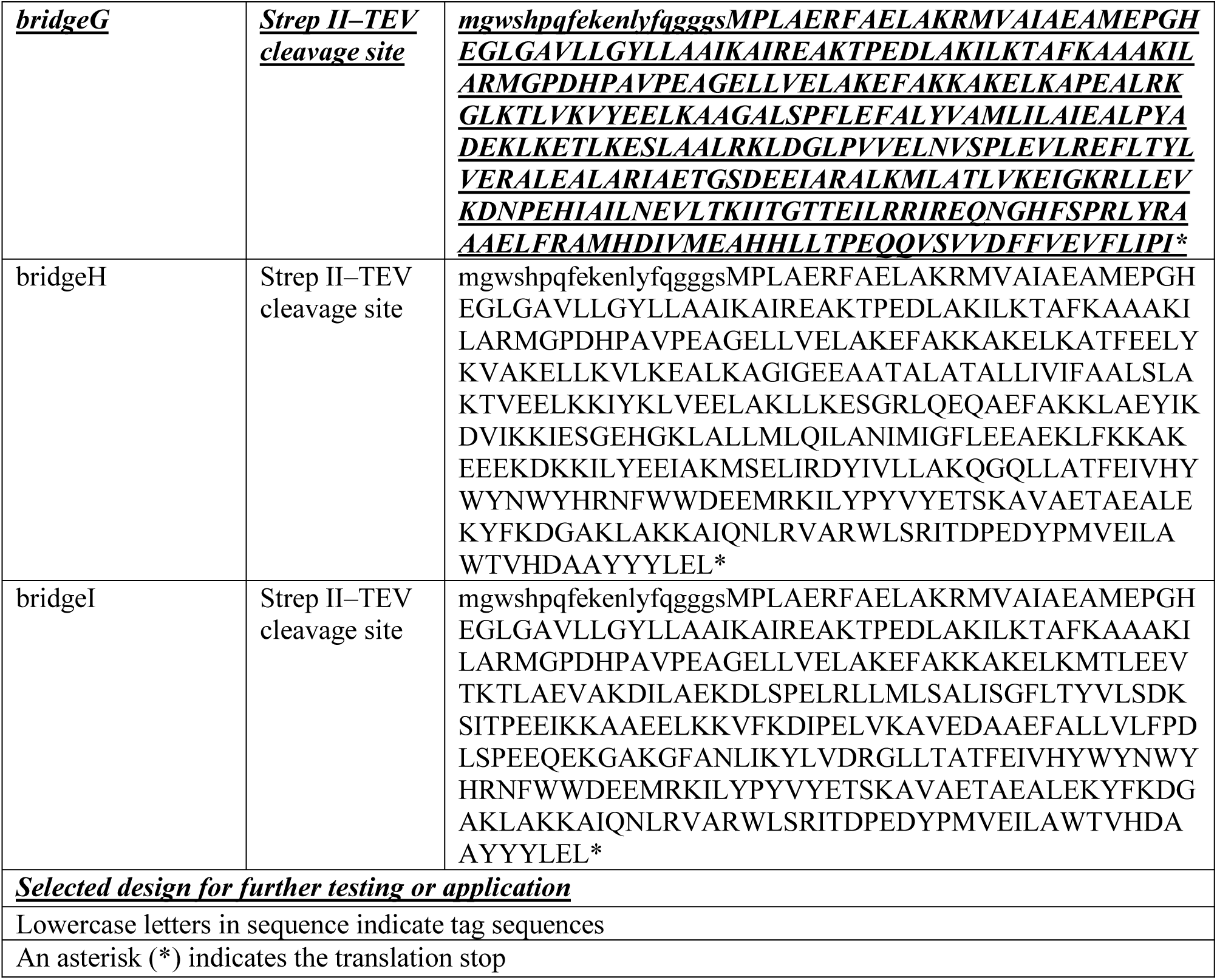
Sequence of designs.

## Materials and Methods

### Shape potential

The shape potential consists of two complementary components: a boundary term and a shape-matching term. The target blueprint was represented as a closed triangular mesh, and a reference point cloud was uniformly sampled from the volume enclosed by the mesh. For the Cα point cloud at the current reverse-diffusion step, 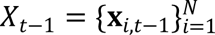, were also treated as a point cloud. For notational simplicity, this point cloud is hereafter denoted 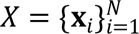. The boundary term was based on the signed distance *d*_SDF_(**x**, ∂Ω), defined as the Euclidean distance to the nearest point on the mesh surface, with positive values assigned to points outside the enclosed volume and negative values to points inside. Only Cα atoms outside the prescribed shape envelope were penalized:

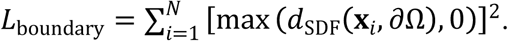

Here, Ω denotes the volume enclosed by the blueprint mesh and ∂Ω denotes its surface. The shape-matching term was defined as the bidirectional Chamfer distance between the current Cα point cloud *X*and the reference point cloud 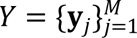 sampled from the volume enclosed by the blueprint mesh:

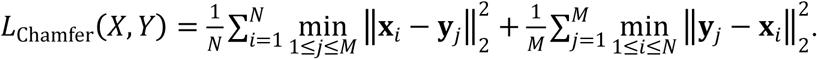

The complete shape potential was defined as

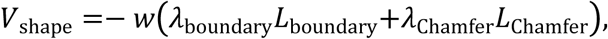

where *w* is the overall guidance weight, and *λ*_boundary_ and *λ*_Chamfer_ determine the relative contributions of the two terms. During diffusion-based backbone generation, the boundary term promotes compliance with the prescribed shape envelope, whereas the Chamfer term promotes global agreement between the evolving backbone and the blueprint volume.

### Generation of blueprints and usage

Blueprints used in this study were generated in OBJ format using Blender (Blender Foundation), although equivalent models could also be created using other CAD software or obtained from publicly available repositories. ChimeraX was used to inspect, scale, and spatially position the blueprint models to ensure consistency with the prescribed shape constraints and intended architectural organization (*34*).

### *De novo* shape-conditioned protein design

Five blueprint-defined target shape envelopes, Cone, L, Sphere, Ellipsoid, and Cube, were selected for proof-of-concept *de novo* shape-conditioned protein generation. All blueprints were centered at the origin in Blender to align with the initial noise distribution and were subsequently verified in ChimeraX.

Protein backbones were generated using RFdiffusion guided by the shape potential. Specifically, 100 backbones were generated for the Sphere and Ellipsoid targets, and 200 backbones were generated for the Cone, L, and Cube targets. Backbone lengths ranged from 200 to 400 residues for the Cone, L, Sphere, and Ellipsoid designs and from 300 to 600 residues for the Cube designs. Because blueprints of different sizes require appropriately sized initial noise representations, we screened backbone lengths within these ranges, and additionally introduced a scale factor to adjust the overall blueprint dimensions. A scale factor of 1.0 preserved the input blueprint coordinates, whereas values of 2.0 and 0.5 uniformly scaled the blueprint coordinates by twofold and one-half, respectively.

Representative backbones that showed close geometric agreement with the prescribed shape envelopes were selected for sequence optimization using ProteinMPNN (*35*). Because the final structure of (*pX0*) was not directly modulated by the shape potential, final backbone geometry was evaluated directly rather than inferred from shape-potential convergence alone. For each selected backbone, eight sequences were initially generated at a sampling temperature of T = 0.3 and evaluated using ColabFold2 (*36*, *37*). Designs with predicted local distance difference test (pLDDT) scores greater than 80 and Cα RMSD values less than 1.5 Å between the predicted and designed backbone models were subjected to a second round of sequence optimization, in which 16 sequences were generated per backbone at T = 0.15. Final designs with pLDDT > 90 and Cα RMSD < 1.5 Å were retained for downstream analysis and experimental characterization.

### Backbone designability analysis

Sequence design was performed using ProteinMPNN followed by structure prediction using ColabFold2, or ESMFold (*38*) depending on the design stage. Predicted structures were evaluated based on model confidence metrics and agreement between predicted and designed backbone conformations using Cα RMSD measurements.

For backbone designability analysis, eight sequences were generated for each backbone using ProteinMPNN with default settings. ESMFold was used for structure prediction. Designs with Cα RMSD < 2.0 Å between predicted and designed backbone structures were classified as sequence-backbone compatible.

### Protein expression and purification of *de novo* shape designs

Synthetic genes encoding the designed proteins with N-terminal His₆-tags in the pET-28a(+) vector (NdeI and XhoI sites) were purchased from Twist Bioscience. Plasmids were transformed into Escherichia coli BL21(DE3) cells and plated on LB agar supplemented with 50 µg/mL kanamycin. Single colonies were cultured overnight in 5 mL LB medium containing kanamycin at 37°C and used to inoculate 1 L LB medium. Cultures were grown at 37°C to an OD₆₀₀ of 0.4–0.6, induced with 0.2 mM IPTG, and incubated at 25°C overnight for protein expression.

Cells were harvested by centrifugation at 5,000 × g for 15 min, resuspended in lysis buffer containing 50 mM Tris-HCl, pH 7.5, 300 mM NaCl, and 25 mM imidazole, and lysed using B-PER reagent (Thermo Scientific). Lysates were clarified by centrifugation at 40,000 × g for 50 min, and the supernatant was applied to Ni-NTA resin (Thermo Scientific) for affinity purification. The resin was washed with buffer containing 50 mM imidazole, and bound proteins were eluted with 300 mM imidazole. Proteins were further purified by size-exclusion chromatography using a Superdex 75 Increase column equilibrated in phosphate-buffered saline (PBS; 137 mM NaCl, 2.7 mM KCl, 10 mM Na₂HPO₄, and 1.8 mM KH₂PO₄, pH 7.4), and monomeric fractions were collected.

### Circular dichroism spectroscopy and thermal denaturation

Circular dichroism (CD) spectra were recorded using a Jasco J-710 spectrometer. Protein samples were prepared at 2–5 µM in PBS, and concentrations were determined by absorbance at 280 nm using a NanoDrop spectrophotometer. Measurements were performed in a 1 mm path-length cuvette at 25°C. Thermal denaturation profiles were monitored at 220 nm while increasing the temperature from 25°C to 95°C at a rate of 1°C/min.

### Crystallization

SC1 crystals were obtained at room temperature by the sitting-drop vapor diffusion method with drops composed of 0.2 µL protein solution (3.4 mg/mL in 20mM Tris-HCl pH 7.8, 40 mM NaCl) and 0.2 µL reservoir solution, which was composed of 1 M tri-Na citrate, 0.1 M Na cacodylate pH 6.5. Crystals were harvested directly from the drop without additional cryoprotection prior to flash-freezing in liquid nitrogen.

SC4 crystals were obtained at room temperature by the sitting-drop vapor diffusion method with drops composed of 0.2 µL protein solution (15.0 mg/ml in 20 mM Tris-HCl pH 7.8, 40 mM NaCl) and 0.2 µL reservoir solution, which was composed of 0.1 M MMT pH 7.0, 25 % w/v PEG 1500. Crystals for both proteins appeared within 48-72 h. Crystals were cryoprotected in mother liquor supplemented with 20% (v/v) ethylene glycol and flash-cooled in liquid nitrogen.

### X-ray diffraction data collection

Diffraction data for SC1 were collected at SSRL beamline BL12-1 using a DECTRIS EIGER 2XE 16M detector (crystal-to-detector distance: 200 mm and a wavelength: 0.97946 Å). A total rotation range of 360° was collected using 0.2° oscillation and 0.2 s exposure per image.

Diffraction data for SC4 were collected at ALS beamline 8.3.1 using a Dectris Pilatus3 S 6M detector (crystal-to-detector distance: 300 mm and a wavelength: 1.11583 Å). A total rotation range of 360° was collected using 0.1° oscillation and 0.1 s exposure per image.

### X-ray diffraction data processing

Diffraction images were integrated with DIALS (*39*) within the xia2 pipeline (*40*). Structures were phased by DIMPLE (*41*) using the spherically-processed data and SC1 or SC4 models as the search models. As the data showed substantial diffraction anisotropy (STARANISO(*42*) reported principal diffraction limits of 2.92, 1.90 and 2.23 Å along a*, b* and c*), the unmerged reflections were reprocessed using autoPROC (*43*) and scaled with AIMLESS (*44*) before an ellipsoidal resolution cutoff was applied using STARANISO based on a local, direction-dependent weighted CC1/2 threshold of 0.30, corresponding to a nominal resolution of 2.30 Å (space group P2₁2₁2₁; unit-cell dimensions a = 60.99, b = 142.63, c = 143.12 Å). Free-R flags were retained from the original xia2/DIALS processing (3.60% of reflections) for all reflections shared between the two datasets to maintain pairwise consistency of the free set across the spherical and anisotropically-truncated data. Flags for the 140 reflections included only with anisotropic resolution extension were generated using GEMMI (*45*) to preserve the same free-reflection fraction and thin-resolution-shell numbering scheme as the original assignment.

### Structure refinement

The structures of SC1 and SC4 were refined against anisotropically truncated data using phenix.refine (*46*), converging to R_work/R_free values of 0.237/0.306 and 0.229/0.270, respectively. The SC1 asymmetric unit contained four protein molecules comprising 1,178 residues and 8,846 non-hydrogen atoms, whereas the SC4 asymmetric unit contained a single protein molecule comprising 355 residues and 2,762 non-hydrogen atoms. Model geometry was validated using MolProbity (*47*). Data collection and refinement statistics are provided in Table S1.

### Topology-preserving shape reshaping

Initial α/β backbone decoys with defined topologies were constructed in Coot, followed by fold-conditioned diffusion with loop insertions of 0–5 residues to diversify backbone length and architecture. A total of 200 trajectories were generated, and backbones exhibiting an α/β two-layer organization were retained for subsequent reshaping analysis. Selected backbones underwent initial sequence optimization using ProteinMPNN and were filtered using pLDDT > 90 and Cα RMSD < 1.5 Å.

To generate left-handed structural states, CAD blueprints encoding left-handed twist were generated in Blender and spatially aligned in ChimeraX relative to the evolving backbone trajectory. This alignment ensured that the initial noise remained contained within the target shape envelope and followed the prescribed growth direction. To sample diverse alignments between the starting backbone and the target shape state, seven blueprint variants were generated by incremental 10° rotational sampling in ChimeraX.

Structure reshaping trajectories were generated for each blueprint-backbone pair. Resulting structures were evaluated based on their agreement with the prescribed left-handed shape state. For each selected backbone, 16 sequences were generated using ProteinMPNN at T = 0.3 and evaluated using ColabFold2. Designs with pLDDT > 80 and Cα RMSD < 1.5 Å were subjected to a second round of sequence optimization, in which 32 sequences were generated per backbone at T = 0.15. Six final designs with pLDDT > 90 and Cα RMSD < 1.5 Å were selected for experimental characterization.

### Protein expression and purification of structure-reshaping designs

Synthetic genes encoding the designed proteins with N-terminal His₆-tags in the pET-28a(+) vector using the NdeI and XhoI sites were purchased from Twist Bioscience. Plasmids were transformed into *Escherichia coli* Rosetta(DE3) cells (Novagen) and plated on LB agar supplemented with 50 µg/mL kanamycin. Single colonies were cultured overnight in 5 mL LB medium containing kanamycin at 37°C and then used to inoculate 1 L Terrific Broth, TB. Cultures were grown at 37°C to an OD₆₀₀ of 0.4–0.6, induced with 0.2 mM IPTG, and incubated overnight at 25°C for protein expression.

Cells were harvested by centrifugation at 5,000 × g for 15 min, resuspended in lysis buffer containing 50 mM Tris-HCl, pH 7.5, 300 mM NaCl, and 25 mM imidazole, and lysed using B-PER reagent (Thermo Scientific). Lysates were clarified by centrifugation at 40,000 × g for 50 min, and the supernatant was applied to Ni-NTA resin for affinity purification. The resin was washed with buffer containing 50 mM imidazole, and bound proteins were eluted with 300 mM imidazole. Proteins were further purified using a HiTrap™ Q HP anion-exchange chromatography column (Cytiva), followed by size-exclusion chromatography using a Superdex 75 Increase column equilibrated in buffer containing 50 mM Tris-HCl, pH 7.4, and 300 mM NaCl. Target-containing fractions were identified by SDS-PAGE and collected for subsequent analyses. For CD experiments, the final size-exclusion chromatography step was performed in PBS, and CD measurements were carried out as described above.

### Cryo-EM structure determination

Purified protein was applied to Quantifoil R1.2/1.3 Cu 400-mesh grids and vitrified using a Vitrobot Mark IV. Grids were screened using a 200 kV Glacios microscope, and grids exhibiting optimal ice thickness and particle distribution were selected for data collection.

For reshaping designs, standard cryo-EM preprocessing, particle classification, and three-dimensional reconstruction workflows were performed using cryoSPARC 4.5 and RELION 5.0 (*48*, *49*). Cleaned particle populations corresponding to monomeric and dimeric states were retained for subsequent three-dimensional reconstruction and refinement. Initial models were generated using ab initio reconstruction. Three-dimensional refinement and local resolution estimation were performed using standard cryo-EM workflows. Final reconstructions were compared with the AF2-predicted design model to assess agreement in global shape, curvature, and secondary-structure organization. Because the available density did not resolve sufficient stereochemical features, such as side-chain density or α-helical pitch, to determine absolute handedness independently, reconstruction handedness was assigned by fitting the stereochemically constrained L-amino-acid AF2 model. Thus, the absolute handedness assignment was model-based, whereas the cryo-EM reconstructions were used to assess structural consistency with the designed left-handed twist state. Data-processing workflows are illustrated in Fig. S4.

For assembly designs, particle picking, two-dimensional classification, and three-dimensional reconstruction were performed using standard workflows with C1 symmetry throughout; no point-group or helical symmetry was imposed. The absolute handedness of each helical reconstruction was similarly assigned relative to the corresponding design model, and the reconstructions were evaluated for consistency with the intended superhelical handedness rather than treated as an independent determination of absolute handedness. A user-defined sharpening B factor of −100 Å² was applied to the final map of hh2-90-4, selected conservatively to improve density visibility without introducing apparent high-frequency artifacts. Structural visualization and analysis were performed using UCSF ChimeraX.

### Design of closed protein architectures

An idealized 340-residue 14-helix bundle was used as a shared starting scaffold for closed assembly design. Three blueprint-defined curvature states corresponding to 90°, 180°, and 270° bends were used to reshape the scaffold into quarter-circle, half-circle, and three-quarter-circle building blocks, respectively.

Building blocks were spatially arranged according to the prescribed architectural organization to generate intermolecular interfaces compatible with closed assembly formation. For the quarter-circle and half-circle architectures, building blocks were rotated according to the target assembly geometry to generate homodimeric backbone models. For the three-quarter-circle architecture, additional in-plane translational adjustment was introduced to facilitate intermolecular interface formation.

For the 180° and 270° architectures, sequence design was performed using two rounds of ProteinMPNN sampling at T = 0.3 and T = 0.15. Symmetry constraints were applied by tying corresponding residues between protomers based on AlphaFold2-predicted homodimeric backbone organization. Designs were filtered using AlphaFold2, AlphaFold3, and Rosetta-based structural metrics.

For the 90° architecture, monomeric backbones were divided into interface and bridging regions. Residues within the bridging regions were constrained between protomers during ProteinMPNN optimization. Interface regions were filtered using AlphaFold2 and Rosetta-based metrics before recombination into full-length constructs (*50*). Final architectures were evaluated using AlphaFold2 monomer prediction and AlphaFold3 homooligomer prediction. Four representative designs per architecture were selected for experimental characterization. The overall sequence design workflow is summarized in Fig. S5.

### Design of open helical architectures

Open helical architectures were generated from the same shared helix-bundle scaffold by introducing local shape states incorporating both curvature and axial twist. Left-handed helical building blocks with local curvature-and-twist states corresponding to 90°, 180°, and 270° helical architectures were generated using structure reshaping. Corresponding right-handed variants were generated using the same reshaping strategy.

Building blocks were spatially arranged to generate intermolecular interfaces compatible with extensible helical assembly formation. All helical building blocks were assembled into homodimeric backbone models prior to sequence optimization. Sequence optimization and structural validation were performed using the same strategy described for the 90° closed architecture. Four representative designs per architecture were selected for experimental characterization. The overall design workflow is summarized in Fig. S5.

### Protein expression of architecture designs

Synthetic genes encoding the designed proteins with N-terminal His₆-SUMO tags in the pET-28a(+) vector using the NdeI and XhoI sites were purchased from Twist Bioscience. Plasmids were transformed into *Escherichia coli* BL21(DE3) cells and expressed as described above. Following Ni²⁺-affinity chromatography, assembly formation was further evaluated by negative-stain transmission electron microscopy.

For selected constructs used in subsequent Cryo-EM and single-particle tracking assay, the His₆-SUMO tag was replaced with a Strep-tag. Cells were harvested by centrifugation at 5,000 × g for 15 min, resuspended in lysis buffer containing 100 mM Tris-HCl, pH 8.0, and 150 mM NaCl, and lysed using B-PER reagent (Thermo Scientific). Lysates were clarified by centrifugation at 40,000 × g for 50 min, and the supernatant was applied to Strep-Tactin®XT 4Flow® resin (IBA) for affinity purification. The resin was washed with lysis buffer, and bound proteins were eluted with elution buffer consisting of lysis buffer supplemented with 50 mM D-biotin, adjusted to pH 8.0. Purified protein was stored at −80 °C until use.

### Negative-stain electron microscopy

Carbon-coated 400-mesh copper grids (Electron Microscopy Sciences) and 1% (w/v) uranyl acetate were used for negative staining following established protocols. Grids were examined using a Tecnai T12 transmission electron microscope (Thermo Fisher Scientific) operated at 120 kV and equipped with a 4k × 4k Gatan Rio 16 CMOS camera.

Micrographs were processed using cryoSPARC. Particle picking, two-dimensional classification, and three-dimensional reconstruction were performed using standard workflows. Representative class averages and reconstructions were compared with the corresponding design models to evaluate agreement with the prescribed architectural organization and external shape states.

### Fluid simulation

Fluid dynamics around the designed assemblies were simulated using custom Python scripts implemented in the Stokes-flow regime using the method of regularized Stokeslets (*24*).

All architectures analyzed in Fig. 5 and Fig. S7 were imported as triangulated surface meshes (in .STL format), including the helical propeller series, symmetric ring controls, and F-actin filament control (PDB: 8F8T). For the helical propeller series, axial lengths were adjusted to be comparable, allowing differences in pumping behavior to be interpreted primarily in terms of helical architecture rather than overall propeller length. Each surface mesh was represented using area-weighted triangular face-center collocation points, with meshes downsampled to a maximum of 2,400 surface elements when required while preserving total surface area.

The surrounding fluid was modeled as water with constant viscosity. Each structure was subjected to prescribed rigid-body rotation with angular velocity Ω = 100 rad s⁻¹. Under the no-slip boundary condition, the surface velocity was defined as u = Ω × r, where *r* denotes the position vector relative to the rotational center. Surface tractions were solved through a regularized-Stokeslet boundary-collocation formulation of the Stokes equations, with the regularization length scale defined from the median surface-element spacing.

Rotation axes were assigned according to the global architecture of each assembly. Slender helical propellers were rotated about the longest principal-component axis. Ring-like assemblies were rotated about the shortest principal-component axis corresponding to the normal of the ring plane. The same rotational axes were used consistently for hydrodynamic simulation, velocity-field visualization, and helical-geometry analysis.

Velocity fields were reconstructed from the solved surface tractions. Axial pumping was quantified using the finite-disk sectional flux

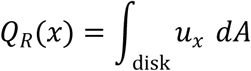

where *u_x_*is the velocity component parallel to the rotational axis and *d* denotes the differential area element on a finite circular disk perpendicular to that axis. Accordingly, *Q_R_*(*x*)represents a local finite-disk axial flux rather than a conserved flux through an infinite cross-section.

For equal-length helical propeller comparisons, axial flux profiles were sampled over a common absolute axial range, and downstream pumping was quantified by averaging *Q_R_*(*x*)within a fixed downstream wake window located 50–200 Å beyond the positive axial end of each assembly. Flux values were normalized by the imposed angular velocity and reported as *Q_R_*(*x*)/Ω.

Helical geometry was measured relative to the same rotational axis used in the hydrodynamic simulations. For helical architectures, an effective helical slope descriptor was defined as *P*/ (*πD*), where *P*is the helical pitch and *D*is the characteristic propeller diameter. This parameter was used to compare pumping trends within the helical architecture series in the context of classical low-Reynolds-number helical propulsion theory.

### F₁-ATPase purification

Two gene fragments encoding the α, β, and γ subunits of PS3 F₁-ATPase from Bacillus sp. PS3 were synthesized with appropriate overhangs by Twist Bioscience. The encoded subunits corresponded to ATP synthase subunit alpha, ATP synthase subunit beta, and ATP synthase gamma chain in PDB 6N2Y, which are annotated as uncA/atpA, uncD/atpD, and uncG/atpG, respectively. The fragments were cloned into the pTrc99a vector (NovoPro Bioscience) by Gibson assembly. The plasmid was further modified as described previously (*52*, *53*), and the N-terminal 18 residues of the α subunit were removed. F₁-ATPase was expressed in the ATP synthase-deficient *Escherichia coli* strain DK8, purification, and functional assays of F₁-ATPase were performed as previously described (*54*). Purified F₁-ATPase was stored in 100 mM potassium phosphate buffer (pH 7.0) at −80 °C until use in assembly tests and single-particle tracking assay.

### Positioning of helical propellers relative to F₁-ATPase

The spatial positioning of the designed helical propellers relative to F₁-ATPase was performed in ChimeraX. The central axis of each helical propeller was aligned with the rotational axis of F₁-ATPase. Multiple candidate propeller-rotor models were generated by rotating the propellers around their central axis. Models with orientations compatible with bridge-mediated binding were selected for subsequent design procedures (Fig. S9A).

### Binder on γ design

Binders targeting the γ subunit of F₁-ATPase were designed using BindCraft (*55*). The γ subunit structure from PS3 F₁-ATPase (PDB: 6N2Y) was truncated and used as the target (Fig. S8A). Two rounds of binder design were performed. In round 1 (R1), binder lengths ranged from 70 to 130 residues, whereas in round 2 (R2), the permitted length range was expanded to 70–200 residues. One hundred designs were generated in each round. The top 10 designs from each round, ranked by BindCraft, were structurally refitted into the positioned propeller-rotor models for orientation evaluation. Four binders from each round were selected for experimental validation (Fig. S8B).

Plasmids encoding the designed binders were synthesized by Twist Bioscience in the pET-28a(+) vector using the NdeI and XhoI cloning sites. Each construct contained an N-terminal Strep-tag. The plasmids were transformed into *E. coli* BL21(DE3). Cells were grown in TB medium supplemented with 50 µg/mL kanamycin at 37°C to an OD600 of 0.4–0.6. Protein expression was induced with 0.2 mM IPTG and continued overnight at 25°C.

Cells were harvested by centrifugation at 5,000 × g for 15 min, resuspended in lysis buffer containing 100 mM Tris-HCl, pH 8.0, and 150 mM NaCl, and lysed using B-PER reagent (Thermo Scientific). Lysates were clarified by centrifugation at 40,000 × g for 50 min, and the supernatant was applied to Strep-Tactin®XT 4Flow® resin (IBA) for affinity purification. The resin was washed with lysis buffer, and bound proteins were eluted with elution buffer consisting of lysis buffer supplemented with 50 mM D-biotin, adjusted to pH 8.0. Proteins were further purified by size-exclusion chromatography using a Superdex 75 Increase column equilibrated in 100 mM potassium phosphate buffer, pH 7.0, as shown in Fig. S8C. Target-containing fractions were assessed by SDS-PAGE.

For binding assays, 400 µg of binder was incubated with 200 µg of F₁-ATPase at room temperature for 30 min. The mixture was then analyzed by size-exclusion chromatography on a Superdex 200 Increase column equilibrated in 100 mM potassium phosphate buffer (pH 7.0) (Fig. S8D). Fractions corresponding to the expected size of the F₁-ATPase complex were analyzed by SDS-PAGE to confirm complex formation (Fig. S8E).

### Bridge design

Successful binders were used to identify binding motifs on the γ subunit. These motifs were reintroduced into the positioned propeller-rotor models. Optimal configurations were selected based on their orientational compatibility for fusion with interface motifs from the helical propeller. Binding motifs and interface motifs were combined with a truncated γ subunit model and used as input for motif scaffolding in RFdiffusion (Fig. S9A). For each motif pair, 100 backbone models were generated. Sequence design was performed using two rounds of ProteinMPNN. Designs were filtered using the following criteria: predicted local distance difference test score (pLDDT) greater than 90, as determined by ColabFold2 in single-sequence mode; and Cα RMSD less than 1.5 Å between the designed and predicted structures. Designs that maintained the proper spatial orientation after re-evaluation within the full assembly model were selected as candidates (Fig. S9B).

Plasmids encoding the selected bridge designs were synthesized by Twist Bioscience in the pET-28a(+) vector using the NdeI and XhoI cloning sites. Each construct contained an N-terminal Strep-tag and a TEV protease cleavage site. The plasmids were transformed into *E. coli* BL21(DE3).

Cells were harvested by centrifugation at 5,000 × g for 15 min, resuspended in lysis buffer containing 100 mM Tris-HCl, pH 8.0, and 150 mM NaCl, and lysed using B-PER reagent (Thermo Scientific). Lysates were clarified by centrifugation at 40,000 × g for 50 min, and the supernatant was applied to Strep-Tactin®XT 4Flow® resin (IBA) for affinity purification. The resin was washed on column with lysis buffer, and bound proteins were eluted by TEV protease cleavage at a protease-to-protein mass ratio of 1:50. Eluted proteins were analyzed by SDS–PAGE (Fig. S9C) and stored at −80 °C until use in assembly tests and single-particle tracking assays.

For binding assays, 20 µg of F₁-ATPase was incubated with bridge protein at a 1:1.5 molar ratio on ice for 15 min. The mixture was purified using Ni-NTA affinity chromatography, followed by buffer exchange into 100 mM potassium phosphate buffer (pH 7.0), using a 100 kDa molecular-weight-cutoff concentrator. A 10-fold molar excess of helical propeller in strep elution buffer was then added and incubated on ice for 2 h. Complexes were enriched using sequential Ni-NTA and Strep-tag affinity purification (Fig. S9D). Final samples were analyzed by negative-stain transmission electron microscopy, nsTEM (Fig. S9E).

### Fluorescent labeling

Strep-tagged hh2-90-4 containing the K271C mutation was purified as described above, except that the lysis buffer supplemented with 2 mM TCEP was used as the wash buffer. The purified protein was stored at −80 °C. After thawing, ATTO 643, prepared as a 10 mM stock solution in DMSO, was diluted in lysis buffer and added to the protein to a final concentration of 1 mM. The labeling reaction was incubated on ice for 2 h. Unreacted dye was subsequently removed using a Zeba Spin Desalting Column (0.5 mL, 7K MWCO; Thermo Fisher Scientific) equilibrated with lysis buffer. The fluorescently labeled protein was subsequently used for assembly in single-particle tracking experiments and for cryo-EM structure determination.

### Fluorescence single-particle tracking assay

For single-particle tracking experiments, purified F₁-ATPase, bridge protein, and fluorescently labeled helical propeller prepared as described above were used. 20 µg F₁-ATPase was incubated with bridge protein at a 1:1.5 molar ratio on ice for 15 min, followed by addition of purified helical propeller at a 10-fold molar excess relative to F₁-ATPase and further incubation on ice for 2 h. The Rotor-Bridge-Propeller complex fraction was enriched by sequential Ni-NTA and Strep-tag pull-down. No buffer exchange using a concentrator was performed during the assembly process. Final samples were collected in Strep elution buffer without EDTA and diluted to a final concentration of approximately 400 pM for single-particle tracking experiments. After dilution, the final buffer contained 25 mM MOPS, pH 7.2, 50 mM KCl, 10% (w/v) tetra-PEG-azide, 2 mM MgCl₂, 5 units/mL pyruvate kinase, and 2 mM phosphoenolpyruvate (PEP). Bridge-Propeller control sample was prepared by mixing the corresponding components at the same initial molar ratios as used for complex assembly and then diluting the samples to achieve comparable fluorescence intensity.

For single-particle tracking experiments, 80 µL of the prepared sample were loaded into a Lab-Tek II Chambered Coverglass System fitted with a #1.5 German coverglass (Thermo Fisher Scientific). Imaging was performed on a Nikon ECLIPSE Ti2 inverted microscope equipped with a 100×/1.49 NA oil-immersion objective and a Hamamatsu ORCA-Flash4.0 scientific CMOS camera (C11440 series). ATTO 643 was excited using a 640-nm laser operated at 15% of the instrument output setting. The excitation beam was directed through the objective, and the incidence angle was adjusted to just above the critical angle for total internal reflection, thereby confining excitation to the coverslip–sample interface and minimizing fluorescence background from the bulk solution. Samples were placed in the sample chamber under temperature control (37°C), and baseline videos were first recorded before ATP addition, referred to as *video^Baseline^* (Fig. 5F). Imaging was initiated after the temperature stabilized at approximately 37°C. ∼1.6 µl 100 mM ATP stock solution prepared in water was then added to the same well to a final concentration of 2 mM. After approximately 15 min, once both the temperature and reaction system had stabilized, a second set of videos was recorded using similar imaging parameters, referred to as *video^ATP^*. Videos were acquired with a frame interval of 50 ms for 500 frames per video. At least three videos containing hundreds of tracks were collected for each condition. The statistics of the single-particle tracking dataset are shown in Table S3. Raw fluorescence time-lapse images were first denoised using DeepCAD (*56*) to reduce shot noise while preserving particle localization and temporal continuity. The denoised TIFF stacks were then analyzed using custom Python scripts. Images were loaded as T × Y × X stacks with a pixel size of 0.16 μm and a frame interval of 0.05 s. For each frame, a Gaussian-filtered background (σ = 14 pixels) was subtracted. Particles were segmented using a robust threshold defined as the image median plus four times the MAD-based noise estimate. Connected components with areas of 4-300 pixels and corrected integrated intensity above 200 were retained. Particle positions were determined from intensity-weighted centroids. Large puncta and artifacts were excluded based on component radius, bounding-box size, and a separate bright-artifact mask. Local background, signal-to-noise ratio, integrated intensity, and aperture-based quality-control metrics were calculated for each detection. Detections were linked into trajectories using Trackpy with a maximum displacement of 1.6 μm, a memory of four frames, and a minimum track length of five localizations. Nearly immobile trajectories were removed if their maximum displacement from the median position was ≤ 0.30 μm and their net displacement was ≤ 0.35 μm.

Stepwise apparent diffusion coefficients were calculated as

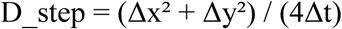

For each trajectory, the arithmetic mean of all valid D_step values was calculated to obtain a trajectory-level mean apparent diffusion coefficient. The trajectory-level values were subsequently averaged within each condition of each independent experiment to obtain the experiment-level mean values used for statistical comparisons. Detection results, trajectories, motion metrics, quality-control plots, and track-overlay visualizations were exported for downstream analysis.

